# Defective plasticity in dermatomyositis patients muscle stem cells is associated with sustained intrinsic inflammatory signaling and disruption of the histone H3.3 chromatin loading pathway

**DOI:** 10.1101/2025.06.09.658580

**Authors:** Wilhelm Bouchereau, Linda Chenane, Lola Lessard, Michèle Weiss-Gayet, Yseult Cardona, Rémi Mounier, Laure Gallay, Yves Allenbach, Olivier Benvéniste, Armelle Corpet, Bénédicte Chazaud, Patrick Lomonte

## Abstract

Skeletal muscle regeneration is driven by muscle stem cells (MuSCs), which proliferate, differentiate and fuse to reform myofibers and restore muscle function. This myogenesis process is driven both by intrinsic MuSC properties and extrinsic cues. While coordinated inflammatory signals are necessary for healthy regeneration, chronic inflammation participates in various pathologies affecting the skeletal muscle. In the idiopathic inflammatory myopathy dermatomyositis (DM), MuSCs exhibit impaired myogenesis *in vitro*, indicating that they may have acquired intrinsic defects, contributing to the disease and providing a mechanism for sustained patient muscle weakness despite efficient anti-inflammatory treatments. Here, we investigated the transcriptomic regulation of DM-derived MuSCs, with a focus on the H3.3 histone variant which regulates myogenesis progression. DM-derived MuSCs were unable to effectively execute the myogenic transcriptional program during in vitro differentiation. They exhibited an activated canonical TNF-⍺ signaling. They also showed reduced expression of H3.3 and its chaperone genes, coupled with a decrease in H3.3 deposition across the entire genome, and particularly at myogenic regulatory factor loci. The loss of H3.3 combined with elevated TNF-⍺ signaling was associated with a failure of DM-derived MuSCs to achieve myogenesis, suggesting a mechanistic link between epigenetic dysregulation and defective muscle regeneration in humans.

## Introduction

Adult muscle stem cells (MuSCs) sustain skeletal muscle regeneration after myofiber damage. They implement adult myogenesis encompassing proliferation, differentiation into myocytes and fusion to form new functional myofibers. Myogenesis is an intrinsic property of MuSCs that is tightly regulated by their environment (Theret and Chazaud, 2024). Muscle regeneration is associated with an inflammatory response where a pro-inflammatory environment initially stimulates MuSC proliferation and represses early differentiation while at later stages, an anti-inflammatory environment promotes differentiation and fusion of MuSCs (Bernard et al., 2022). Sustained inflammation is linked to muscle wasting and MuSC dysfunction in ageing (Moiseeva et al., 2023). However, the impact of chronic inflammation on MuSC biology has not been thoroughly investigated. Dermatomyositis (DM) is a rare disease belonging to the family of idiopathic inflammatory myopathies. It is characterized by high levels of type I interferon (IFN-I), which correlates with the severity of the disease (Bolko et al., 2021; Greenberg et al., 2012). While anti-inflammatory treatments such as glucocorticoid administration improve patient’s health, many of them experience muscle weakness (Landon-Cardinal et al., 2020; Pinal-Fernandez et al., 2017). This autoimmune disease is characterized by permanent myofiber damage, associated with a strong inflammatory environment due to the presence of a huge leukocyte infiltrate within the skeletal muscle. We have previously shown that MuSCs from DM patient’s muscle (DM-MuSC) proliferate and differentiate much less than healthy control (HC)-MuSCs (Gallay et al., 2022). Those intrinsic defects of myogenesis likely participate in the disease by impairing muscle regeneration. DM-MuSCs offers the opportunity to investigate the intrinsic effects of long-term inflammation on MuSCs.

Cell identity is maintained by several layers of epigenetic regulations. These mechanisms preserve cell specificity over time while allowing a degree of plasticity to respond to environmental cues. Controlled deposition and recycling of histone variants is such a mechanism. Studies in various mouse cell types have shown that the replication-independent deposition of the H3.3 histone variant is crucial for safeguarding cell identity and is necessary for the proper regulation of myogenic regulatory factor (MRF) transcription and myogenesis (Esteves de Lima et al., 2021; Fang et al., 2018; Funk et al., 2022; Saleh et al., 2018). Also, chromodomain-helicase-DNA-binding protein 2 (CHD2)-dependent deposition of H3.3 allows proper myogenic differentiation, while the repression of either CHD2 or H3.3 leads to the loss of MRF expression and to a defective myogenic differentiation (Harada et al., 2012; Yang et al., 2011). Finaly, the loss of H3.3 chaperone protein Histone Cell Cycle Regulator (HIRA) leads to the downregulated expression of skeletal muscle genes and to the upregulated expression of genes from other lineages (Esteves de Lima et al., 2021). Despite compelling evidence from mouse models highlighting the importance of H3.3 deposition in regulating myogenesis and maintaining cell identity, these mechanisms remain largely uncharacterized in human systems. DM offers a unique pathological context to explore the impact of chronic inflammation on H3.3 dynamics and its consequences for human MuSCs function. Notably, H3.3 deposition is known to be altered in inflammatory contexts. In senescent cells, both HIRA and H3.3 relocalize to promoters of active genes in line with transcriptional changes (Rai et al., 2014), and during viral infection or interferon stimulation, increased HIRA recruitment enhances H3.3 deposition at interferon-stimulated genes (ISGs) (Kleijwegt et al., 2023; McFarlane et al., 2019). These findings suggest that chronic inflammatory signals in dermatomyositis may similarly reshape H3.3 dynamics, thereby contributing to defective myogenesis.

In this study, the transcriptomic profiles of human primary DM-MuSCs and HC-MuSCs were analyzed throughout the myogenesis process, which can be recapitulated in vitro. DM-MuSCs displayed markedly reduced transcriptomic plasticity from proliferation to differentiation. The role of IFN-I signaling in this defect was found to be limited. In contrast, several inflammatory pathways, particularly those associated with Tumor Necrosis Factor (TNF)-α, were found to be overexpressed in DM-MuSCs. At the chromatin level, a widespread reduction in H3.3 deposition was observed. Notably, treatment of HC-MuSCs with TNF-α recapitulated both the myogenesis defects and the global decrease in H3.3 deposition seen in DM-MuSCs.

## Results

### DM-MuSCs exhibit widespread suppression of the myogenic program and an enhanced inflammatory transcriptomic signature

We previously showed that DM-MuSCs present intrinsic myogenesis defects (Gallay et al., 2022). As a follow up, an RNA-seq analysis was conducted on those cells to compare with that of HC-MuSCs. Proliferation refers to cells cultured in growth medium (Prolif), early differentiation occurs 1 day after transitioning to differentiation medium (DiffD1), and late differentiation occurs 3 days after the shift to differentiation medium (DiffD3). Differential gene expression analysis of DM-*vs.* HC-MuSCs was performed in each culture condition. Gene set enrichment analyses (GSEA) of the differentially expressed genes (DEGs) show that in all conditions, DM-MuSCs overexpress genes involved in inflammatory processes including TNF-⍺, Interleukin (IL)-6 or IFN signaling (Fig. 1A-left, Supplementary Table 1). They also exhibit a significant downregulation of genes involved in myogenesis, consistently identified as the most significantly downregulated term in each GSEA analysis (Fig. 1A-right, Supplementary Table 1). DM-MuSCs in Prolif and DiffD1 conditions also show a downregulation of the genes associated with proliferation and cell-cycle progression such as G2M checkpoints and E2F targets (Fig. 1A-right, Supplementary Table 1). This is coherent with DM-MuSCs having a lower proliferation rate than HC-MuSCs (Gallay et al., 2022). In contrast, they show a higher expression of those genes at DiffD3 (Fig. 1A-left, Supplementary Table 1), suggesting a delay in cell cycle exit upon differentiation. DEGs between DM and HC-MuSCs show little overlap between the 3 conditions along the myogenesis process. Twenty-five genes (out of 749) show common upregulation, including pro-inflammatory genes (*HLA-DRB1*, *IFI44*) and pro-fibrotic genes (*FN1*) (Fig. 1B). Expression of only 21 genes is commonly decreased in all states including genes involved in development (*HOTAIR, HOX*) (Fig. 1B). The myogenesis process is known to have a drastic impact on both transcriptome and epigenome (Long et al., 2022; Massenet et al., 2023; McKellar et al., 2021). Principal component analysis (PCA) reveals that the primary transcriptomic variations across samples is driven by the differentiation process rather than by the origin from DM or HC muscle (Supplementary Fig. S1A). To go further, we looked at the kinetics of expression of DEGs between DM- and HC-MuSCs along the differentiation process. Genes which expression is upregulated in DM-MuSCs at Prolif or DiffD3 stages remain highly expressed as compared with HC-MuSCs during the whole myogenic process (Fig. 1C left, Supplementary Fig. S1B). Conversely, genes which expression is downregulated in DM-MuSCs remain weakly expressed as compared to HC-MuSCs during all the myogenic process (Fig. 1C right, Supplementary Fig. S1B). Overall, although the DM-MuSC transcriptome follows the myogenic program similarly to that of HC-MuSCs, there is a significant global expression shift between DM and HC-MuSCs, suggesting that DM-MuSCs have a slower dynamic and a reduced capacity to timely remodel their transcriptome during myogenesis.

**Figure 1.**
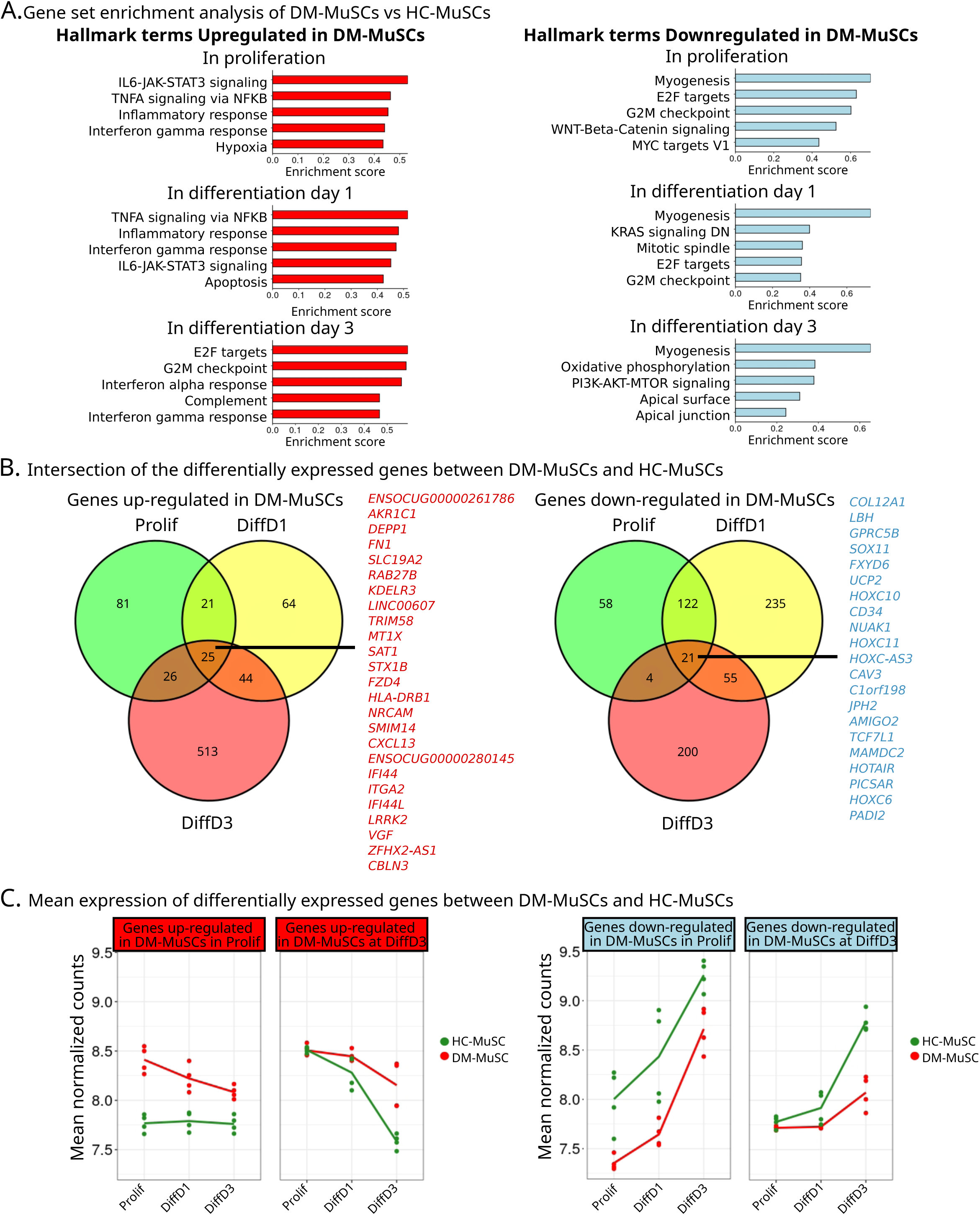
Comparison of HC- and DM-MuSC transcriptome. (**A**) Molecular Signatures Database Hallmark (MSigDB Hallmarks) gene set enrichment analysis (GSEA) results on the differential expression analysis between DM- and HC-MuSCs in the 3 conditions, ranked by enrichment scores. Left panels show Hallmark terms enriched in DM-MuSCs, right panels show Hallmark terms under-represented in DM-MuSCs. (**B**) Overlap of the differentially expressed genes (DEGs) in Prolif, DiffD1 and DiffD3 conditions between DM- and HC-MuSCs. The lists of the common up-regulated (left panel, in red) and down-regulated (right panel, in blue) DEGs in DM-MuSCs are given. (**C**) Mean expression in DM- and HC-MuSCs at the 3 time points (Prolif, Diff1, Diff3) of the DEGs that were identified by comparing DM- and HC-MuSC gene expression at Prolif (green circles in (**B)** and Supplementary Fig. 1B) and at DiffD3 (red circles in (**B)** and Supplementary Fig. 1B) with upregulated DEGs presented in the left panel (red) and downregulated DEGs presented in the right panel (blue)s.

### DM-MuSCs show a reduced transcriptomic plasticity

The impact of chronic inflammatory state on the transcriptomic dynamics of DM-MuSCs in proliferation and differentiation was then evaluated. As standard control, DEG analyses in HC-MuSCs along the 3 steps of myogenesis was first performed (Supplementary Fig. S2 and Supplementary Table 4). Genes associated with chemotaxis and hypoxia are over-represented in Prolif as compared with DiffD1 (Supplementary Fig. S2Aa). As expected, genes related to muscle development and differentiation are more enriched at DiffD1 as compared with Prolif (Supplementary Fig. S2Ab). In the subsequent step of myogenesis, i.e., DiffD1 vs. DiffD3, terms associated with DNA replication and mitotic division are over-represented in DiffD1 as compared with DiffD3 (Supplementary Fig. S2Ba). Inversely, and as expected, an enrichment is observed for genes associated with muscle function, development and differentiation, at DiffD3 as compared with DiffD1 (Supplementary Fig. S2Bb). These results are coherent with what is expected for the *in vitro* myogenesis process. We next compared in HC- and DM-MuSCs the average expression of genes in the 4 DEG sets described above (Supplementary Table 2). Genes that are more expressed in HC-MuSCs in Prolif *vs.* DiffD1 show a decreased expression along the differentiation process in both HC- and DM-MuSCs, though they are more expressed in the latter. Those genes are notably related to chemotaxis (for instance *CXCL2)* (Fig. 2A and S2Aa). The expression of DEGs upregulated in DiffD1 *vs*. DiffD3 in HC-MuSCs decreases exponentially along differentiation, as exemplified by genes involved in cell-cycle progression such as *CCNB1* (Fig. 2B and S2Ab). Importantly, DM-MuSCs fail to properly repress this DEG set (Fig. 2B). This suggests a reduced capacity of DM-MuSCs to exit the cell-cycle during differentiation. Finally, DEGs upregulated in DiffD1 *vs*. Prolif (Supplementary Fig. S2Ab) and in DiffD3 *vs*. DiffD1 (Supplementary Fig. S2Bb) show a sustained increased expression during the entire differentiation process (Fig. 2C and 2D). They show an enrichment in genes related to muscle differentiation and development such as *MYMX* or *MYH3* (Fig. 2C and 2D). Interestingly, both DEG sets maintain a lower expression level in DM-MuSCs *vs.* HC-MuSCs (Fig. 2C and 2D), suggesting a delay for DM-MuSCs in entering differentiation. Overall, these results mark a reduced transcriptomic plasticity of DM-as compared with HC-MuSCs throughout differentiation, with a delay in exiting the cycle cycle and entering into terminal myogenic differentiation.

**Figure 2.**
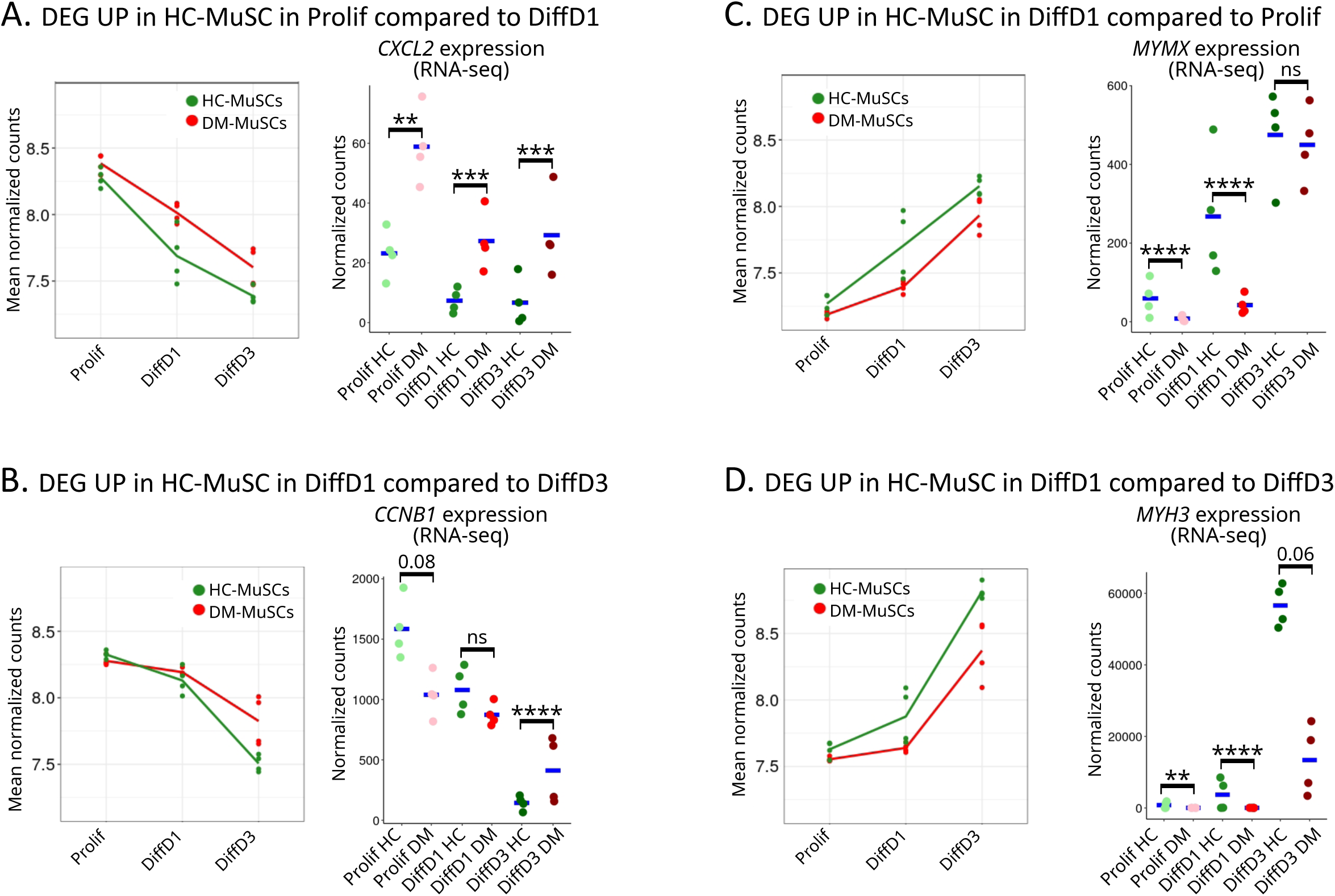
DM-MuSCs show reduced plasticity in response to differentiation stimulus. In the left panels, the mean expression of DEGs identified in Supplementary Fig. S2 (HC-MuSCs along the myogenesis process) was evaluated in both HC (green)- and DM (red)-MuSCs in the 3 conditions. The right panels show the expression in the RNA-seq data of selected genes belonging to the corresponding DEGs, with p-value significance between HC- and DM-MuSC indicated on the graph. (**A**) DEGs upregulated in Prolif (Prolif *vs*. DiffD1). Expression of *CXCL2*. (**B**) DEGs upregulated in DiffD1 (DiffD1 *vs*. DiffD3). Expression of *CCNB1*. (C) DEGs upregulated in DiffD1 (Prolif *vs*. DiffD1). Expression of *MYMX*. (**D**) DEGs upregulated in DiffD3 (DiffD1 *vs*. DiffD3). Expression of *MYH3*.

This blunted plasticity of DM-MuSCs is confirmed in the enrichment maps generated from the over-represented terms in HC and DM-MuSCs in the DiffD3 condition (Supplementary Fig. S3). As expected, a strong enrichment of terms associated with muscle development and contraction is observed in HC-MuSCs (Supplementary Fig. S3A), whereas DM-MuSCs remain abnormally enriched in terms associated with proliferation and cell cycle (Supplementary Fig. S3B). Hence, cell cycle regulators, such as cyclins and cyclin-dependent kinases, are more strongly expressed in HC-MuSCs as compared with DM-MuSCs during proliferation (Supplementary Fig. S3C Prolif, and S3D). They sustain a quicker repression in HC-MuSCs as compared with DM-MuSCs as the differentiation process progresses to DiffD1 then DiffD3 (Supplementary Fig. S3C DiffD1 and DiffD3, and S3D, Supplementary Table 3). For instance, the proliferation marker MKI67 shows a marked tendency toward higher expression in HC-MuSCs as compared with DM-MuSCs under proliferative conditions, a pattern that appears reversed in the DiffD3 condition (Supplementary Fig. S3E). This was confirmed at the protein level, MKI67+ cells are more abundant in HC- as compared with DM-proliferating MuSCs (Supplementary Fig. S3F), while conversely, DM-MuSCs retain a higher number of MKI67+ cells than HC-MuSCs in the DiffD3 condition (Supplementary Fig. S3G). Similar results were obtained using EdU (Supplementary Fig. S3H-I). A specific focus was then made on genes related to myogenesis. As expected, the expression of most of the genes associated with myogenesis and muscle functions, including myosin heavy chains coding genes, *RYR1, MYMX* and *MYMK,* show a strong increase in the DiffD3 condition in HC-MuSCs (Fig. 3A and B). Although their expression kinetics follow a similar increase in DM-MuSCs, their initial expression level is lower, and they reach those observed in HC-MuSCs (Fig. 3A and B). The initial low expression was confirmed by RT-qPCR of 6 myogenic genes in proliferating cells and shows that all but *MYOD1* have a lower expression in DM- as compared with HC-MuSCs (Supplementary Fig. S3J). The deficit in myogenic differentiation of DM-MuSCs is confirmed by the lower expression of *MYOG* mRNAs in DM-*vs*. HC MuSCs (Fig. 3C) and a lower number of DM-MuSCs expressing MYOG protein as compared with HC-MuSCs (Fig. 3D) in the DiffD3 condition. Taken together, these results indicate that DM-MuSCs fail to suppress proliferation-associated genes during differentiation, to upregulate MRFs, and exhibit an intrinsic impairment in rewiring their myogenic program.

**Figure 3.**
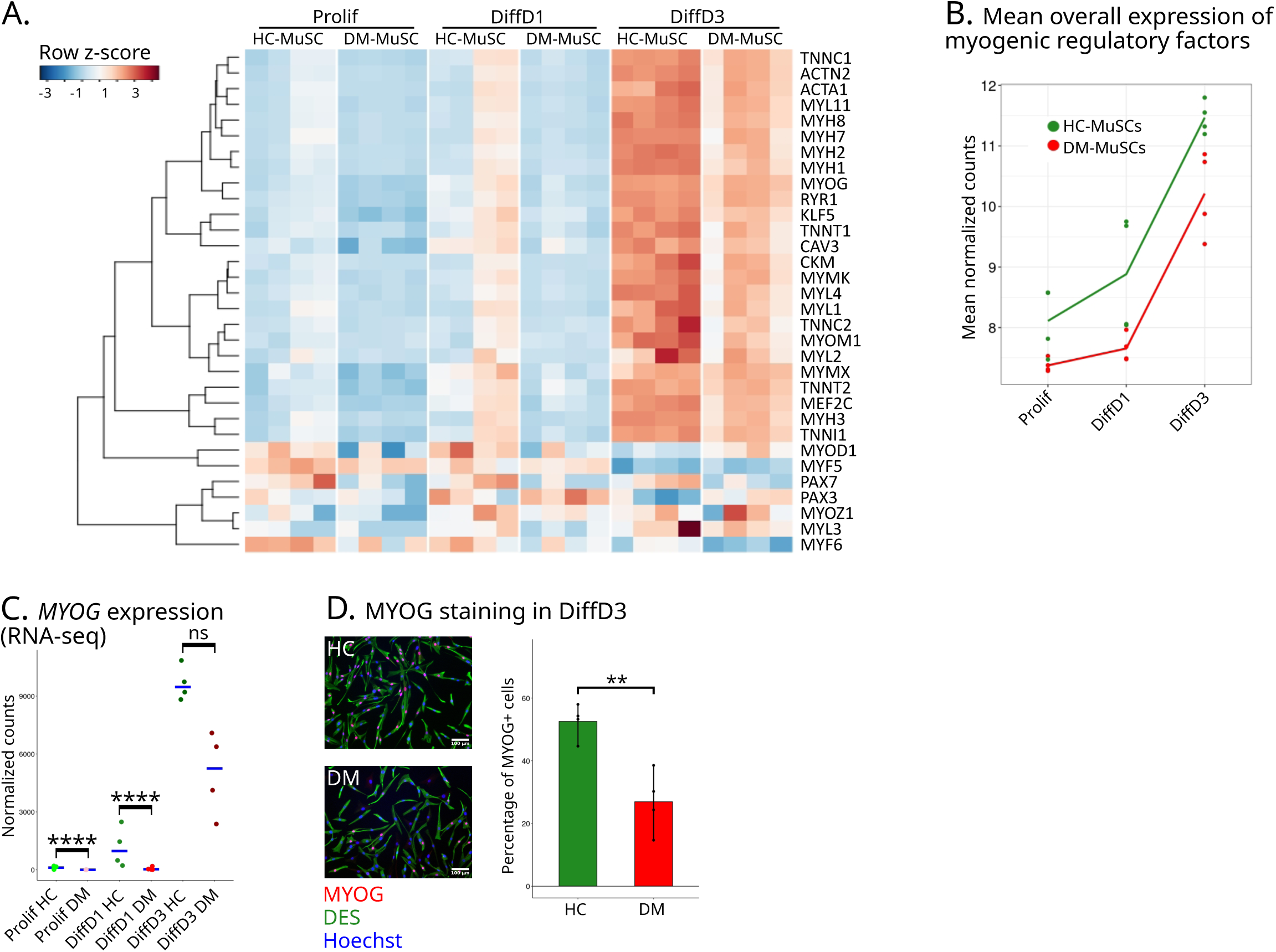
DM-MuSCs show an impaired myogenesis transcriptional program. (A) Heatmap of the expression of myogenic regulatory factors and their targets in DM- and HC- MuSCs in Prolif, DiffD1 and DiffD3 conditions. (**B**) Mean expression of myogenic regulatory factors and their targets in the 3 conditions by DM- (red) and HC- (green) MuSCs. (**C**) Expression of MYOG from the RNA-seq., blue crossbars indicate median for each cell type in each condition. P-value significance between HC- and DM-MuSC is indicated on the graph. (**D**) Immunofluorescence for MYOG (red) in HC- and DM-MuSCs, from four different patients each. Cells are labelled for desmin (green) and nuclei with hoechst (blue) and quantification of the percentage of MYOG+ nuclei in each sample. P-value significance between HC- and DM-MuSC is indicated on the graph.

### DM-MuSCs show an intrinsic activation of pro-inflammatory signaling pathways

Pro-inflammatory stimulation alters MuSC differentiation (Saclier et al., 2013). The above transcriptomic data show that DM-MuSCs overexpress genes involved in several pro-inflammatory pathways (Fig. 1A). The expression of 84 genes including receptors, ligands and targets of IFN type I and II, IL-1, IL-6 and TNF-⍺ pathways was investigated. Overall, these genes are highly expressed in DM- as compared with HC-MuSCs at all stages of the myogenesis process, with their expression decreasing along the course of myogenesis (Fig. 4A and B and Supplementary Fig. S4A). Notably, DM-MuSCs express higher levels of chemokines and Major Histocompatibility Complex (MHC) coding genes than HC-MuSCs (Supplementary Fig. S4B). This tendency was confirmed on a subset of genes by RT-qPCR (Supplementary Fig. S4C). Strikingly, the expression levels of the two ISGs, *MX1* and *ISG15*, remain comparable between DM-MuSCs and HC-MuSCs in Prolif (Fig. 4C and S4C), even though they are known to be over-expressed in DM skeletal muscle (Allenbach et al., 2016; Amici et al., 2021; Salajegheh et al., 2010). DM is characterized by a strong IFN signature, present both systemically in leukocytes and locally in skeletal muscle, with a correlation to disease severity (Amici et al., 2021; Argyriou et al., 2023; Gallay et al., 2019; Greenberg et al., 2012). However, when the expression of 48 ISGs (McFarlane et al., 2019) was analyzed, only a few of them show significant overexpression in DM-MuSCs (labelled in red in Fig. 4C, Supplementary Table 4). These results suggest that overall pro-inflammatory pathways are maintained active in DM-MuSC. This does not seem to be the case with IFN signaling pathway, in spite of its well described involvement in DM.

**Figure 4.**
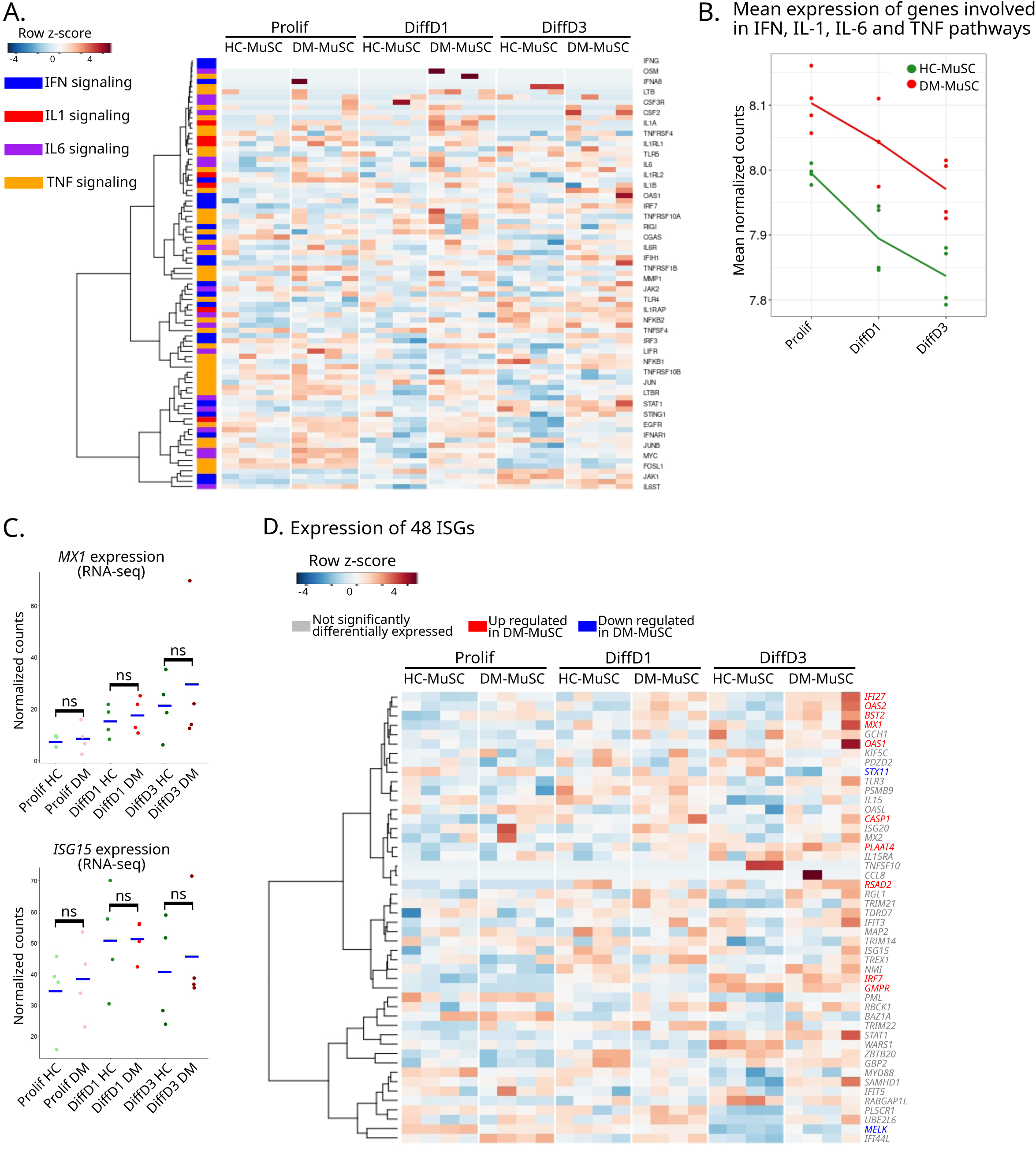
DM-MuSCs show intrinsic activation of pro-inflammatory pathways. (**A**) Heatmap of the expression of all the genes involved in type I, II and III IFN signaling (blue), IL1 signaling (red), IL6 signaling (purple) and TNF-α (orange) family signaling between DM- and HC-MuSCs in Prolif, DiffD1 and DiffD3 conditions. (**B**) Mean expression of all the genes involved in type I, II and III IFN signaling, IL1 signaling, IL6 signaling and TNF-⍺ family signaling between DM- (red) and HC-MuSCs (green) in Prolif, DiffD1 and DiffD3 conditions. (**C**) Expression of *MX1* and *ISG15* from the RNA-seq data. Blue crossbars indicate median for each cell type in each condition. P-value significance between HC- and DM-MuSC is indicated on the graph. (**D**) Heatmap of the expression of 48 ISGs from (McFarlane et al., 2019) between DM- and HC-MuSCs in Prolif, DiffD1 and DiffD3 conditions. Gene names are coloured in red and blue if they are significantly up- and down-regulated between DM- and HC-MuSCs, respectively.

To further investigate IFN signaling, MuSCs were treated with IFN-β to analyze if they could reactivate a memory IFN-I response (Kamada et al., 2018). IFN-β treatment has no effect on proliferation and commitment into differentiation of both DM-MuSCs and HC-MuSCs (Supplementary Fig. S5A). Accordingly, IFN-β treatment of HC-MuSCs does not impact on their capacity to express myogenic genes (but induced the expression of its targets *MX1* and *OAS1*) both in Prolif and DiffD3 conditions (Supplementary Fig. S5B and S5C). It is known from previous studies that IFN-β treatment represses high density-cultured HC-MuSC fusion (Franzi et al., 2013; Ladislau et al., 2018). We successfully replicated these findings, confirming that HC-MuSCs used in our study responded as expected (Supplementary Fig. S5D). Taken together, these data indicate that IFN signaling alone has a limited impact on the early stages of myogenesis in normal MuSCs in culture, suggesting that other signaling pathways may be responsible for the impaired plasticity observed in MuSCs from DM patients.

### Activation of pro-inflammatory pathways induces loss of myogenesis

As detailed above, DM-MuSCs exhibit a much higher expression of genes associated with various inflammatory pathways as compared with HC-MuSCs (Fig. 4A and S4A). Common transcription factor (TF) signature on the promoters of the DEGs revealed that in all 3 conditions (Prolif, DiffD1 and DiffD3) genes that are underexpressed in DM-MuSCs are targets of myogenic TFs (*e.g.*, MEF2, MYF5, MYOD, MYOG) (Fig. 5A, Supplementary Table 5). In contrast, overexpressed genes are enriched in TF motifs of NFkB-p65-Rel and the AP-1 family (e.g., JUNB, BATF, FOS, FOSL2, FRA1, FRA2), which are canonical targets of the TNF-⍺ signaling pathway. Additionally, an increase of the Cell Cycle Homology Region (CHR) motifs is specifically found in DM-MuSCs in the DiffD3 condition (Fig. 5A, Supplementary Table 4). CHR motifs are linked to the transcriptional regulation of cell cycle genes, consistent with the elevated expression of proliferation-associated genes in DM-MuSCs (Supplementary Fig. S3A,C,D). These findings suggest that an intrinsic activation of the canonical TNF-⍺ signaling pathway is at work in DM-MuSCs during the differentiation process. We therefore analyzed molecular and cellular parameters of HC-MuSCs treated with TNF-α in comparison with IFN-β. Similarly to IFN-β, no effect of TNF-α, in combination or not with IFN-β is observed on HC-MuSC proliferation (Fig. 5B). Interestingly, unlike IFN-β, TNF-α strongly repress MuSC differentiation, as monitored by MYOG+ cells, and without synergizing with IFN-β (Fig. 5C). Accordingly, TNF-α treatment strongly up-regulates the expression of its targets (*TNFAIP3* and *IL6)* while also increasing that of IFN (*MX1* and *OAS1*) in proliferating (Fig. 5D) and differentiating (Fig. 5E) conditions. While the stimulation of ISGs by TNF-α may not be entirely unexpected, it is not a commonly reported finding in the literature. We thus re-analyzed data from a previous study investigating the impact of TNF-α treatment on skeletal myogenesis using murine C2C12 myoblasts (O’Brien et al., 2020), and confirmed the TNF-α-dependent increase in the expression of representative ISGs (Supplementary Fig. S6). Moreover, TNF-α, unlike IFN-I, significantly repress the expression of myogenic regulatory factors *PAX7* and *MYOD1* in proliferating HC-MuSCs (Fig. 5D) and repress the expression of *MYOG* and *MYH3* in differentiating cells (Fig. 5E). To minimize potential biases from acute stimulation, HC-MuSCs were treated for 8/9 days with TNF-α or IFN-β (Supplementary Fig. S7A). The results are similar to those of the acute treatment, showing no impact on proliferation (Supplementary Fig. S7B) and a decrease in differentiation (Supplementary Fig. S7C). The transcriptomic expression of MRFs, IFN and TNF-⍺ targets mirrors those observed with the short treatments (Supplementary Fig. S7D and S7E). Altogether, these results show that IFN-I has only a minimal direct impact on myogenesis while TNF-α stimulation significantly impairs the differentiation of HC-MuSCs. TNF-α also induces the expression of ISGs, thereby mimicking the dysfunctions and altered gene expression observed in DM-MuSCs. This suggests that DM-MuSCs may have a cell-intrinsic activation of TNF-⍺ signaling, which both sustains inflammation and disrupts differentiation.

**Figure 5.**
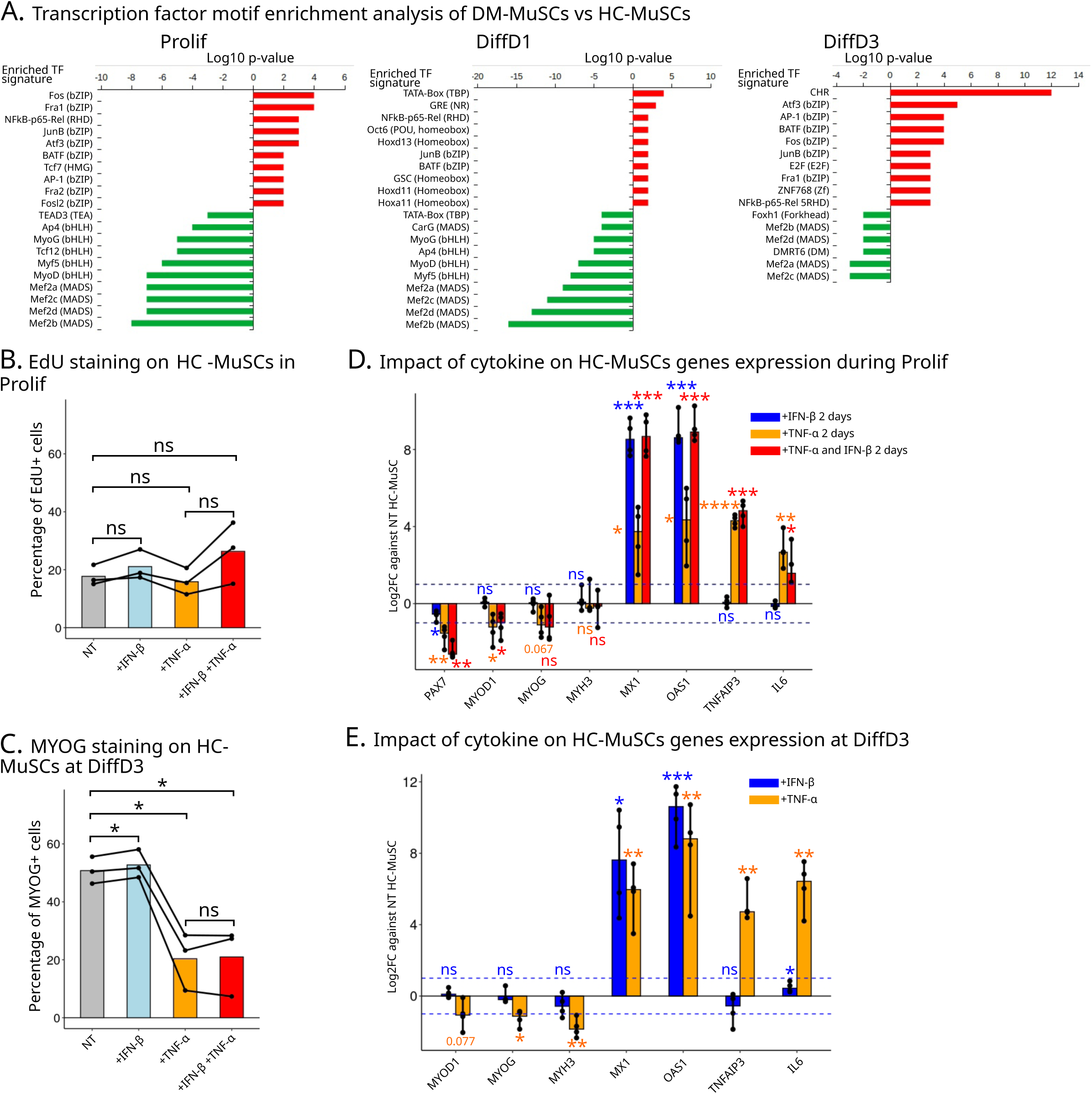
TNF-α stimulation impairs myogenesis. (**A**) Transcription factor (TF) motif analysis of the promoters of DEGs between DM- and HC-MuSCs in Prolif (left panel), DiffD1 (middle panel) and DiffD3 (right panel). In y-axis, the name of the enriched TF motif signatures, with in parenthesis the corresponding DNA-binding domain family. Motifs enriched in promoters of DEGs up- and down-regulated in DM-MuSCs appear in red and green, respectively. (**B**) Quantification of the mean percentage of EdU+ nuclei in HC-MuSCs from three different patients in Prolif condition with or without addition of IFN-β and/or TNF-α. P-value significance between HC- and DM-MuSC is indicated on the graph. (**B**) Quantification of the mean percentage of MYOG+ nuclei in HC-MuSCs from four different patients in DiffD3 condition with or without addition of IFN-β and/or TNF-α. P-value significance between HC- and DM-MuSC is indicated on the graph. (**D**) Expression of myogenic genes, IFN- and TNF- stimulated genes of HC-MuSCs from four different patients treated with or without IFN-β and/or TNF-α in the Prolif condition (RT-qPCR). P-value significance between HC- and DM- MuSC is indicated on the graph. (**E**) Expression of myogenic genes, IFN- and TNF-stimulated genes of HC-MuSCs from four different patients treated with or without IFN-β or TNF-α in the DiffD3 condition. P-value significance between HC- and DM-MuSC is indicated on the graph.

### DM-MuSCs show global histone H3.3 reduction and loss of transcriptomic plasticity

Our data show that DM-MuSCs exhibit severe myogenesis defects and maintain pro-inflammatory pathways active, even when removed from their inflammatory tissue environment and long-term cultured. We thus hypothesized that DM-MuSCs might sustain epigenetic dysregulations, acting as a long-term memory that contributes to their intrinsic myogenesis defects. Strikingly, H3.3 coding genes (*H3F3A* and *H3F3B*) and several of its chaperones are among the significantly repressed DEGs in DM-MuSCs in the Prolif condition (Fig. 6A and S8A) as well as for *H3F3A* in the DiffD3 condition (Supplementary Fig. S8B). Dynamics of the histone variant H3.3 is known to be impacted by inflammatory stresses such as senescence (Rai et al., 2014), viral infection, and IFN stimulation (Kleijwegt et al., 2023; McFarlane et al., 2019). A decrease of H3.3 was confirmed at the protein level using western blot in the Prolif condition (Supplementary Fig. S8C). DM-MuSCs also show a decrease in the expression of essential components of the H3.3 chaperone HIRA complex *UBN1* in the Prolif condition (Fig. 6A) and *ASF1A* in both Prolif and DiffD3 conditions (Fig. 6A and S8B). Hence, the genome-wide deposition of H3.3 in HC- and DM-MuSCs was assessed by CUT&RUN in Prolif and DiffD3 conditions. Differential binding analysis of H3.3 between HC- and DM-MuSCs reveals a general reduction of H3.3 binding at most consensus peaks in DM- as compared with HC-MuSCs in both Prolif and DiffD3 conditions (Fig. 6B and S8D). These results fit with a global loss of H3.3 across the chromatin, as expected from the reduced expression of histone H3.3 and H3.3 chaperones coding genes in DM-MuSCs. We next investigated whether the loss of H3.3 was uniform throughout the chromatin. We looked at the deposition of H3.3 on MRFs, ISGs and inflammation-related genes in HC- as compared with DM-MuSCs. In the Prolif condition, all gene sets have decreased H3.3 binding, while in the DiffD3 condition, H3.3 signal is highly decreased on MRFs, unchanged on ISGs, and moderately decreased on inflammation-related genes in DM-MuSCs as compared with HC-MuSCs (Fig. 6C). Analysis of the CUT&RUN results for the distribution of H3.3 in DM-MuSCs and HC-MuSCs on the DEGs shows that both up-and down-regulated genes in DM-MuSCs incorporated significantly less H3.3 than HC-MuSCs in the Prolif condition (Supplementary Fig. S9A). At DiffD3, the differential incorporation of H3.3 in down-regulated genes in DM-MuSCs is even more exacerbated (Supplementary Fig. S9B). To assess the functional impact of altered H3.3 deposition, we integrated CUT&RUN data with RNA-seq under proliferative conditions. Overall, changes in gene expression and H3.3 chromatin binding between DM- and HC-MuSCs show only a weak correlation (Supplementary Fig. S9C). However, genes with both reduced H3.3 enrichment and decreased expression are significantly enriched for myogenesis-related signatures (Supplementary Fig. S9D). These results suggest that although DM-MuSCs exhibit a global decrease in H3.3 incorporation across the genome, this deficiency is particularly pronounced at MRFs, and becomes even more marked during differentiation. Interestingly, treatment of proliferating HC-MuSCs with TNF-α treatment tends to reduce H3.3 deposition at the transcription end sites (TES) of both myogenic regulators and inflammatory genes (Supplementary Fig. S9E), yet it does not repress the expression of the H3.3-encoding genes, *H3F3A* and *H3F3B* (Supplementary Fig. S9F). These findings suggest that DM-MuSCs have a deficiency in H3.3 deposition on chromatin, resulting from both reduced expression of H3.3 and its chaperones, as well as pro-inflammatory cytokine induced decrease of H3.3 deposition.

**Figure 6.**
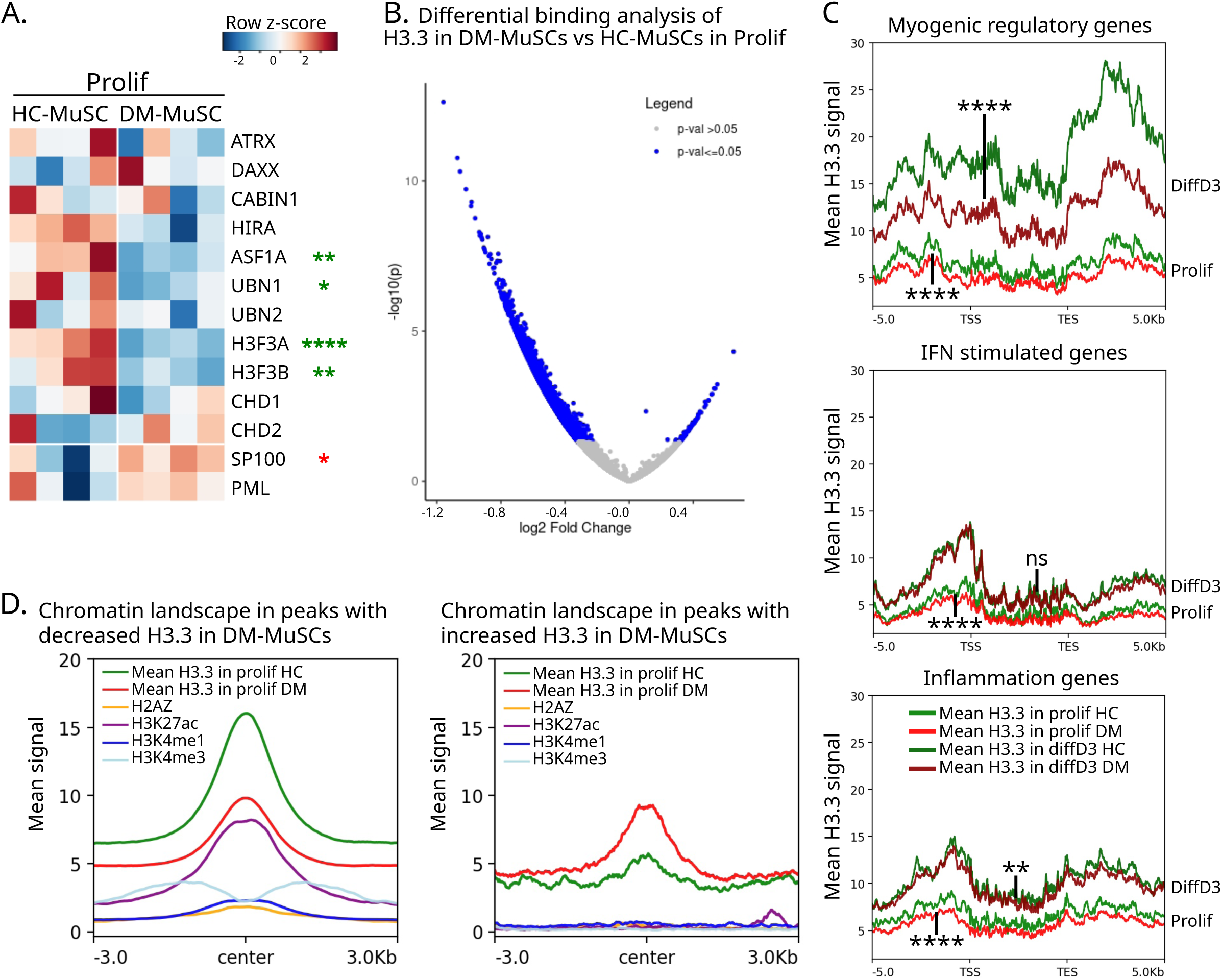
DM-MuSCs have a global loss of H3.3. (**A**) Heatmap of the expression of H3.3 coding genes, H3.3 chaperones and PML and SP100 in HC- and DM-MuSCs in the Prolif condition. Genes significantly down or up-regulated between HC- and DM-MuSCs are labelled with red and green asterisks, respectively (the number of asterisks refers to the P value). (**B**) Volcano plot of the differential binding analysis of H3.3 between DM- and HC-MuSCs in the Prolif condition. Peaks with a significant differential binding of H3.3 appear in blue. (**C**) Intensity plot of the mean H3.3 signal on myogenic regulatory genes (top panel), IFN type I stimulated genes (middle panel) or inflammation-related genes (bottom panel) in DM- (light red) andf HC- (light green) MuSCs in Prolif condition, and in DM- (dark red) and HC- (dark green) MuSCs in DiffD3 condition. Differential mean H3.3 signal within the -5/+5kb of the TSS and TES of the selected genes was evaluated between HC- and DM-MuSCs, with each gene of the selected panel counted as a replicate. P-value significance between HC- and DM- MuSC is indicated on the graph. (**D**) Intensity plot on differentially bounded peaks with decreased (left panel) or increased H3.3 binding (right panel) in DM-MuSCs. Mean H3.3 signal in DM (red) and HC- (green) MuSCs in Prolif condition. Mean signal of H2AZ in yellow, H3K27ac in purple, H3K4me1 in dark blue and H3K4me3 in light blue in HC-MuSCs (datasets from ENCODE, Zhang et al., 2020, https://www.encodeproject.org/).

It is well-established that post-translational modifications of histones can influence their effect on the chromatin environment, with H3K4me3 primarily found at active promoters, H3K4me1 at enhancers, and H3K27ac at both active promoters and enhancers (Pchelintsev et al., 2013). To further explore the correlation between H3.3 loss in DM-MuSCs and the dysregulation of specific genomic regions, we compared existing ENCODE ChIP-seq datasets for histone modifications in Prolif HC-MuSCs (Zhang et al., 2020, https://www.encodeproject.org/) with our results on the genome-wide deposition of H3.3 in DM-MuSCs. Interestingly, H3.3 peaks with decreased signal in DM-MuSCs are more enriched in active chromatin marks H3K27ac, H3K4me1, H3K4me3 and histone variant H2AZ than the DM-MuSCs-related peaks (Fig. 6D). More specifically, about 16.4% of the peaks with increased H3.3 level in HC-MuSCs are shared with peaks of H3K4me1 and H3K27ac (Supplementary Fig. S9G). This would predict that H3.3 is predominantly lost at active enhancers in DM-MuSCs, potentially explaining their reduced capacity to remodel the transcriptome in response to differentiation cues. In conclusion, loss of H3.3 in both MRFs and in regulatory elements could be at the roots of the trancriptomic dysregulation we observe in DM-MuSCs, and lead to their inability to properly undergo myogenesis.

## Discussion

In this study, major transcriptional defects are observed in DM-MuSCs, with a global up-regulation of pro-inflammatory genes and repression of myogenic-associated genes. DM-MuSCs exhibit reduced transcriptional plasticity, failing to properly exit the cell cycle and to activate the myogenic transcriptional program. Although they retain their myogenic identity, they fail to activate key MRFs. DM-MuSCs show an intrinsic activation of canonical TNF-⍺ signaling, with the up-regulation of targets of NFκB and AP-1 family of TFs. Strikingly, this intrinsic inflammation is not marked by intracellular elevated IFN-I signaling, which corroborates with the fact that IFN-I only weakly impacts MuSC dynamics *in vitro* (only the last fusion step). TNF-α stimulation of HC-MuSCs partly mimics DM-MuSCs by directly blunting their differentiation. Additionally, DM-MuSCs exhibit reduced expression of H3.3 and its chaperones, which correlates with lower H3.3 levels on the chromatin. Along with the intrinsic effects of TNF-α, this suggests that defects in H3.3 dynamics could further impair the activity of myogenic enhancers, directly contributing to the dysfunction of MuSCs in DM. However, it remains unclear whether the failure of DM-MuSCs to upregulate MRFs is a consequence of reduced H3.3 deposition or if the impaired upregulation of MRFs itself leads to decreased H3.3 deposition.

DM-MuSCs present an intrinsic elevation of pro-inflammatory signaling. However, while DM is characterized by a high IFN-I signature, DM-MuSCs have a moderate intrinsic IFN-I signaling activity. Given the lack of effect of IFN-I on the first steps of *in vitro* myogenesis observed in this study, this suggests that IFN signaling may not directly influence proliferating or early differentiating MuSCs. Instead, IFN-I stimulation represses cell fusion. Other studies reported a negative effect of IFN-I on MuSC fusion and myofiber size (Franzi et al., 2013; Ladislau et al., 2018), while another publication showed that on a 3D myobundle model, IFN-I stimulation leads to reduced muscle contraction strength and increased fatigue (Covert et al., 2024). Therefore, elevated IFN-I levels may have a greater impact on myofibers than on MuSCs. Another potential implication of elevated IFN-I could be on the skeletal muscle microenvironment. Indeed, another characteristic of DM is the infiltration of immune cells in the perifascicular area, mainly monocytes and neutrophils (Li et al., 2023; Rostasy et al., 2004; Yasin et al., 2019; Zhao et al., 2023). Immune cells are believed to be the main source of IFN-I and other pro-inflammatory cytokines found in DM (Li et al., 2023; Ye et al., 2022). However, the mechanism driving their activation and infiltration into the muscle remains poorly understood. Interestingly, DM-MuSCs up-regulate the expression of chemoattractants, such as CXCL chemokines. There is a growing number of publications reporting beneficial effect of JAK inhibitors on DM patients (Ladislau et al., 2018; Landon-Cardinal et al., 2023; Paik et al., 2021; Zhang et al., 2023). JAK inhibitors suppress IFN-I signaling, potentially protecting myofibers, while also reducing activation of the immune system (Ladislau et al., 2018).

In spite of the reported high IFN-I activity in DM patients, IFN-I signaling pathway appears weakly active in DM-MuSCs *in vitro.* Instead, they show intrinsic activation of the canonical TNF-α signaling pathway. TNF-α superfamily ligands and downstream effectors are well known for their capacity to block differentiation of MuSCs (Chen et al., 2007; Kim et al., 2020; Ogura et al., 2013; Ono and Sakamoto, 2017; Schmidt et al., 2021; Shirakawa et al., 2021). In addition, elevated TNF-α signaling has been described in other context with chronic inflammation, such as sarcopenia (Bian et al., 2017; Dalle et al., 2017), ageing (Li et al., 2020; Wang et al., 2018) and cancer cachexia (Mantovani et al., 2000). Overexpression of TNF-α in mice leads to early onset of rheumatoïd arthritis, loss of muscle mass and interstitial lung disease (Bell et al., 2019), the latter being also common among MDA5+ DM patients (Allenbach et al., 2020). Nonetheless, our results, together with the existing literature, suggest that strong of TNF-α stimulation alone is insufficient to fully replicate the physiological defects of DM-MuSCs, especially since it does not impact on MuSC proliferation (Li, 2003; Shirakawa et al., 2021). While it has been demonstrated that low concentration of TNF repress proliferation, it also increases myogenic differentiation (Chen et al., 2007). This opens at least two possibilities: either an alternative mechanism is responsible for a persistent repression of the proliferation of DM-MuSCs, or long-term, chronic exposure to TNF-α could have broader effects as compared with short-term stimulation. A recent publication further complexifies the role of TNF-α in the regulation of myogenesis (Fu et al., 2024), where TNF-α knock-down in MuSCs increases both proliferation and differentiation. Thus, it seems that autocrine TNF-α signaling in MuSCs inhibits both their proliferation and differentiation, which is highly reminiscent of the phenotype of DM-MuSCs. The differences between endogenous and exogenous TNF-α stimulation in MuSCs remain to be explored, particularly given the very low expression of the *TNFA* gene in both DM-MuSCs and HC-MuSCs.

The up-regulation of inflammatory signaling pathways can partly explain DM-MuSC myogenesis defects. However, the mechanism sustaining their intrinsic inflammation remains elusive. Indeed, how can inflammation be sustained without the presence of the immune cells and its proinflammatory signals? Several recent publications show that following acute inflammation, AP1 family of TFs are retained on some enhancers, associated with the retention of euchromatin marks H3K27ac and H3K4me1 and highly open chromatin (Falvo et al., 2023; Larsen et al., 2021). These loci and the binding of AP1 TF allow an inflammatory memory response following a second stimulation. One study shows that the inflammatory exposure of pancreatic cells leads to improper cell fate decision and maintenance of an inflammatory specific population of de-differentiated cells (Falvo et al., 2023). This is reminiscent of our results that show failure of DM-MuSCs to properly engage in differentiation and their maintenance of an inflammatory phenotype. The authors also showed that this correlates with the retention of H3K4me1 and H3K27ac on inflammation memory loci, with an associated AP1 signature. In our study, we observed an elevated expression of targets of AP1 TF in DM-MuSCs no matter the condition, even though there is a poor conservation of the dysregulated genes from one condition to the other. This could be explained by AP1 targeted enhancers being constantly active in DM-MuSCs, but shifting their target genes during the differentiation process as gene regulatory networks are remodeled. Chronic exposure to inflammation of DM-MuSCs could have led to a pathological memory response, which would maintain an inflammatory specific population of DM-MuSCs with elevated activation of AP1 and NFkB signaling pathways. Inflammatory MuSCs could propagate inflammation to healthy cells or at least prevent their proper differentiation by the repression of differentiation induced by their secretion of TNF-α or other proinflammatory cytokines.

In addition to intrinsic TNF-α activity, DM-MuSCs present blunted response to myogenesis and fail to remodel their transcriptome to properly differentiate. Interestingly, there is a big overlap between MuSC specific enhancers and regions with reduced H3.3 in DM-MuSCs. Deposition of H3.3, notably by HIRA, has been associated with the regulation of cell fate and cell identity (Esteves de Lima et al., 2021; Fang et al., 2018; Funk et al., 2022; Saleh et al., 2018). Knock-out of either HIRA or H3.3 in mouse embryonic stem cells reduces the binding of de novo expressed TFs to their target loci, preventing proper differentiation. This highlights the crucial role of H3.3 deposition for cell fate (Tafessu et al., 2023). Moreover, H3.3 is also enriched in poised enhancers marked by H3K4me1 but no other active marks (Pchelintsev et al., 2013). It was shown in mice that knock-out of HIRA leads to the reduced incorporation of H3.3 in the chromatin and consequently the reduction of active mark H3K27ac at ATAC peaks loci. This ultimately impedes myogenesis by MuSCs (Esteves de Lima et al., 2021). Finally, retinoic acid stimulation followed by time-ChIP revealed that H3.3 is already present on retinoic acid activated enhancers before the onset of the up-regulation of retinoic acid induced genes (Chen et al., 2013). Thus, H3.3 deposition would bookmark poised enhancers and maintain their chromatin open to be rapidly activated upon intrinsic or extrinsic stimulation. In DM-MuSCs, loss of H3.3 could lead to the erosion of the euchromatin on myogenic-specific enhancers, reducing their capacity to be rapidly activated upon differentiation. This could explain the loss of transcriptional plasticity observed in DM-MuSCs. Our study reveals for the first time a potential link between chronic inflammation and loss of H3.3, associated with an intrinsic activation of TNF-α signaling. This opens new avenues to investigate how inflammation can become chronic and pathological in myopathies.

## Materials & methods

### Patient cells selection

Human skeletal MuSCs derived from DM patients were selected from a previous study (Gallay et al., 2022). Standard protocol approvals, registrations, patient consents and case selection methods are detailed in (Gallay et al., 2022). Rapidly, patients without immuno-suppressive and cytotoxic drugs administration before muscle biopsy were selected. In the previous study, histologic staining and immunohistochemistry of muscle sections were performed to evaluate the histologic dermatomyositis severity score. In the current study we chose 4 patients with high severity on this score, which together with other clinical data indicate severe muscle harm. A summary of relevant patients’ information and clinical data can be found in Supplementary Table 6. More detailed information can be found in (Gallay et al., 2022). MuSCs derived from the 4 DM patients were obtained from the Hospices Civils de Lyon cell bank (Cardiobiotec CRB-HCL AC2013-1867). All experiments involving DM-MuSCs were done with those cells. MuSCs from healthy controls paired on sex and age were obtained from surgical waste of patients undergoing programmed knee or hip surgery.

### Cell culture

MuSCs derived from healthy controls were directly extracted by mechanical and enzymatic digestion from fresh muscle biopsies (Gallay et al., 2022). After an expansion period, they were frozen and stored in liquid nitrogen until further use. After thawing, MuSCs were purified at the first passage of primary culture using anti-CD56 coated magnetic beads (Miltenyi Biotech 130-050-401) to obtain pure myogenic cell cultures as described previously (Gallay et al., 2022). Cell purity was checked by flow cytometry (Miltenyi Biotech, MACSQuant Analyzer) using anti-CD56 antibodies (APC-conjugated anti-CD56 antibodies, BD Pharmingen #555518) and isotypic control (BD Pharmingen #555751).

MuSCs were cultured in complete medium (Skeletal Muscle Cell Growth Medium #23060 Promocell, supplemented with Mix C-39365, Promocell), 10% of heat-inactivated Fetal Bovine Serum (FBS, Sigma-Aldrich) and 1% of Penicillin/streptomycin (Gibco, 14190169). Cells were passaged upon reaching 80% confluency using 0.05% trypsin-EDTA (Gibco, 25300-054) digestion then seeded back onto new supports.

### Proliferation assay

Analysis of proliferation of MuSCs was performed as follows: MuSCs were seeded at 3000 cells/cm² on labteks (Dutscher, 177437). The next day, medium was changed with fresh medium. 24 hours later, 50% of the medium was replaced with fresh medium. Proliferation was evaluated at day 2, using 5-ethynyl-29-deoxyuridine (EdU) incorporated for 6 hours (kit EdU Clickit Alexa Fluor 488; Invitrogen). At the end of the proliferation assay, cells were washed with PBS twice and fixed with formaldehyde 4%. Fixed cells were permeabilized cells in PBS containing 0.5% Triton X-100. For proliferation assay, EdU was revealed for 30min according to the manufacturer instructions (kit EdU Clickit Alexa Fluor 488; Invitrogen). Cells were washed twice in PBS and blocking was performed with 1 hour incubation in BSA 4% (diluted in PBS, Fisher scientific, BP9701). Incubation with MKI67 primary antibody (Abcam, ab15580) diluted in BSA 4% was performed overnight. Secondary antibodies were then incubated for 40 min at 37°C (See Supplementary Table 7 for antibodies used). After washing several times with PBS the samples, they were incubated with streptavidin dylight (Vector laboratories, SA-5549) for 20 min at 37°C. Cells were washed thoroughly with PBS, and cells were briefly incubated in Hoechst 33342 (Sigma-Aldrich, 14533) for nuclei visualization. Slides were mounted using Fluoromount G (Invitrogen, 15586276).

### Differentiation Assay

Analysis of differentiation capacities of MuSCs was performed as follows: cells were seeded in labtek at 1500 cells/cm2 in growth medium. The next day, medium was changed with differentiation medium with or without cytokines consisting of skeletal muscle cell basal medium (Promocell, C-23260) supplemented with 10 μg/mL insulin (Sigma-Aldrich, I2643), FBS 0.5% and 1% of Penicillin/streptomycin. 50% of the medium was replaced with fresh medium every day for 3 days. Proliferation was evaluated at day 3 by using 5-ethynyl-29-deoxyuridine (EdU) incorporated for 6 hours (kit EdU Clickit Alexa Fluor 488; Invitrogen). At the end of the proliferation assay, cells were washed with PBS twice and fixed with formaldehyde 4%. Fixed cells were permeabilized cells in PBS containing 0.5% Triton X-100. For proliferation assay, EdU was revealed for 30min according to the manufacturer instructions (kit EdU Clickit Alexa Fluor 488; Invitrogen). Cells were washed twice in PBS and blocking was performed with 1 hour incubation in BSA 4% (diluted in PBS, Fisher scientific, BP9701). Incubation with anti-MYOG (BD Pharmingen, 556358) and anti-DES (Abcam, ab32362) primary antibodies diluted in BSA 4% was performed overnight. Secondary antibodies were then incubated for 40 min at 37°C (See Supplementary Table 7 for antibodies used). After washing several times with PBS the samples, they were incubated with streptavidin dylight (Vector laboratories, SA-5549) for 20 min at 37°C. Cells were washed thoroughly with PBS, and cells were briefly incubated in Hoechst 33342 (Sigma-Aldrich, 14533) for nuclei visualization. Slides were mounted using Fluoromount G (Invitrogen, 15586276).

### High-density differentiation Assay

Analysis of differentiation capacities of MuSCs was performed as follows: cells were seeded in labtek at 10 000 cells/cm2 in growth medium. The next day, medium was changed with differentiation medium consisting of skeletal muscle cell basal medium (Promocell, C-23260) supplemented with 10 μg/mL insulin (Sigma-Aldrich, I2643), FBS 0.5% and 1% of Penicillin/streptomycin. 50% of the medium was replaced with fresh medium every day for 3 days. At the end of the high-density differentiation assay, cells were washed with PBS twice and fixed with formaldehyde 4%. Fixed cells were permeabilized cells in PBS containing 0.5% Triton X-100. Cells were washed twice in PBS and blocking was performed with 1 hour incubation in BSA 4% (diluted in PBS, Fisher scientific, BP9701). Incubation with anti-MYOG (BD Pharmingen, 556358) and anti-DES (Abcam, ab32362) primary antibodies diluted in BSA 4% was performed overnight. After washing several times with PBS the samples, they were incubated with streptavidin dylight (Vector laboratories, SA-5549) for 20 min at 37°C. Cells were washed thoroughly with PBS, and cells were briefly incubated in Hoechst 33342 (Sigma-Aldrich, 14533) for nuclei visualization. Slides were mounted using Fluoromount G (Invitrogen, 15586276).

### Acute cytokine assay

In order to assess the effect of acute treatment with cytokines on proliferation, cells were grown as follows: MuSCs were passaged and seeded at 3000 cells/cm² in growth medium. The next day, medium was replaced with fresh one with cytokines. TNF-α was added at 5 ng/ml (Invivogen, rcyc-htnfa) while IFN-β at 1000 U/ml (Peprotech, 300-02BC). 50% of the medium was changed the next day with fresh growth medium with cytokines. At day 2, cells were either processed for proliferation assay as described above or for RNA-extraction followed by Reverse transcription (RT) – qPCR, or for chromatin immunoprecipitation (ChIP) - qPCR (see below). In order to assess the effect of acute treatment with cytokines on differentiation, cells were grown as follows: MuSCs were passaged and seeded at 1500 cells/cm² in growth medium. The next day, medium was replaced with fresh one with cytokines. TNF-α was added at 5 ng/ml (Invivogen, rcyc-htnfa) while IFN-β at 1000 U/ml (Peprotech, 300-02BC). 50% of the medium was changed the next 2 days with fresh growth medium with cytokines. At day 3, cells were either processed for differentiation assay as described above or for RNA-extraction followed by RT-qPCR (see below).

### Mid-term cytokine assay

In order to assess the effect of mid-term treatment with cytokines on proliferation, cells were grown as follows: MuSCs were passaged and seeded at 3000 cells/cm² in growth medium. The next day, medium was replaced with fresh one with cytokines. TNF-α was added at 5 ng/ml (Invivogen, rcyc-htnfa) while IFN-β at 1000 U/ml (Peprotech, 300-02BC). 50% of the medium was changed every day with fresh growth medium with cytokines. Cells were passaged upon reaching 80% confluency as detailed above. Cells were grown for 5 days before passaging them for either proliferation or differentiation assay as described in the acute cytokine assay protocol (see above).

### Microscopy, imaging, and quantification

Images were acquired with the Axio Observer Z1 inverted wide-field epifluorescence microscope (Zeiss) and a CoolSnap HQ2 camera from Photometrics. Identical settings and contrast were applied for all images of the same experiment to allow data comparison. For each sample, 5-10 pictures were recorded at 10x magnification. Raw images were treated with Fiji software. Proliferation index was defined as the number of EdU-positive or MKI67-positive nuclei over the total number of nuclei. Differentiation index was defined as the number of Myogenin-positive nuclei over the total number of nuclei. For differentiation at high density, fusion index was calculated as the number of nuclei present in myotubes (>2 nuclei in the same desmin-positive cell) over the total number of nuclei and Differentiation index was defined as the number of Myogenin-positive single nuclei added to nuclei present in myotubes over the total number of nuclei.

### Reverse transcription (RT)

At the end of the proliferation or differentiation assay, cells were detached by trypsin-EDTA incubation, washed with PBS and pelleted at 400g for 5min. Pellets were snap-frozen and stored at - 80°C before RNA-extraction. TRIzol reagent protocol (Invitrogen, 15596026) was used to isolate total RNAs, resuspended in ddH2O according to the manufacturer instructions. Contaminant DNA was removed by digestion with DNAse for 15 min at room temperature (Promega, M610A) and reaction was stopped with STOP buffer (Promega, M199A) incubation for 2min at room temperature. We used 500 ng of RNA for RT. RNAs were annealed with Random Primers (Invitrogen, 58875) and RT was performed with the RevertAid H Minus Reverse Transcriptase (Thermo Scientific, FEREP0451) according to the manufacturer instructions. cDNAs were stored at –20 °C before qPCR analysis.

### ChIP

At the end of the proliferation assay, cells were crosslinked directly in the culture dishes as described previously (Cohen et al., 2018). After the PBS washes, cell pellets were snap-frozen in liquid nitrogen and stored at –80 °C before immunoprecipitation. Cells were de-frozen on ice and chromatin was prepared following the TruChIP protocol from Covaris, as described in (Cohen et al., 2018). We used the Covaris M220 Focused-ultrasonicator to shear through chromatin (7 min at 140 W, Duty off 10%, Burst cycles 200). After shearing, chromatin immunoprecipitation was performed. We used 20 μL of protein A magnetic dynabeads (Invitrogen, 10001D) for immunoprecipitation with 2 μg of anti-H3.3 antibody (Diagenode, C15210011). DNA was purified with phenol-chloroform (Sigma-Aldrich, 77617). Samples were centrifuged at 20000g at 4°C for 5 min. Liquid phase was harvested, then 1ug/ml of glycogen, 200mM of NaCl and 2 volumes of cold ethanol 100% was added to each sample. DNA was precipitated for 2 hours at -20°C, then centrifuged for 10 min at 4°C and 20000g. Supernatant was discarded. Pellets were washed with ethanol 70% and centrifuged 5 min at 4°C and 20000g. DNA pellets were dried then resuspended in ddH20 and stored at –20 °C before qPCR analysis.

### Quantitative PCR (qPCR)

QPCRs were performed using the ONE SYBR qPCR Master Mix (Ozyme, OZYA008). Primers used for qPCR are described in the Key Resources Supplementary Table 7 in Appendix.

### Western blot analysis

At the end of the proliferation or differentiation assay, total cellular extracts were obtained by directly lysing the cells in 2 X Laemmli sample buffer (LSB) (125 mM Tris-Hcl pH 6.8, 20% glycerol, 4% SDS, bromophenol blue) containing 100 mM DTT. Lysate was subjected to 15% acrylamide gel electrophoresis, and transfer was then run overnight on nitrocellulose membrane (Thermo Scientific, 88018). Blocking was done by incubating membranes one hour at room temperature with 5% BSA (diluted in PBS). Antibodies were incubated overnight at 4°C or 1 hour at room temperature. Anti-rabbit antibody coupled to HRP was then incubated for 2 hours at room temperature, and the membrane was revealed with either ECL (Thermo Scientific, 32109) or ECL femto (Thermo Scientific, 34096).

### RNA-seq

At the end of the proliferation or differentiation assay, cells were detached by trypsin-EDTA incubation, washed with PBS and pelleted at 400g for 5min. Pellets were snap-frozen and stored at - 80°C before RNA-extraction. QIAGEN Rneasy mini kit was used to extract and purify RNA according to manufacturer instructions (Qiagen, 74104). Purified RNAs were stored at -80° before library preparation and sequencing. NEBNext® Ultra™ II Directional RNA Library kit (New England Biolabs) was used to prepare an RNA-seq library. Paired-end sequencing was performed on an Element AVITI™ system (HELIXIO, Clermont-Ferrand, France).

### RNA-seq analysis

RNA-seq paired-reads were aligned and quantified using salmon (version 1.9.0) on gencode v41 for index generation. Resulting counts were processed and analyzed on R (version 4.4.1) using Deseq2 package (version 1.44.0). DEG were selected with a p-value under 0,05 and a log2FoldChange either above 1 or under -1. Heatmap and lineplots were produced using counts normalized by variance stabilizing transformation. Gene set enrichment analysis was performed using GSEA (version 4.3.2) and using log2FoldChange values as ranking. Over-representation analysis was performed with enrichplot (version 1.24.0), org.Hs.eg.db (3.19.1) and clusterProfiler (4.12.0) packages. Transcription factor motif enrichment analysis was performed on DEG using the findMotifs.pl function of HOMER (version 3.12).

### CUT&RUN

At the end of the proliferation or differentiation assay, 250 000 cells were detached by trypsin-EDTA incubation, washed with growth medium and pelleted at 600g for 3min. Samples were then processed with active motif CUT&RUN kit (Active motif, 53180) following manufacturer instructions with some modifications inspired from (Ding and Braun, 2023). Nuclei were first isolated by resuspending pellets in nuclei isolation buffer and incubating them for 10 min on ice. Nuclei were then washed twice in wash buffer and bound to activated Concavalin A beads for 10 min at room temperature. Following that, nuclei were incubated with antibodies overnight at 4°C with 1 ug of anti-H3.3 antibody (Active motif, 91191). After antibody incubation, unbound antibodies were washed twice with permeabilization buffer and pAG-MNase was added. Samples were incubated for one hour at 4°C. Cells were washed twice with permeabilization buffer then cooled to 0°C and incubated with ice-cold 100 mM CaCl2 for 2 h at 4°C. MNase digestion was terminated with the addition of STOP buffer. RNA and protein digestion was performed by incubating samples overnight at 37°C with 0.1% of SDS and 0.3 mg/ml of proteinase K. DNA was purified by phenol-chloroform as detailed in the ChIP protocol. DNA was stored at –20 °C before library preparation. The libraries were paired-end sequenced on a DNBSEQ Technology platform (BGI Genomics Co., Shenzhen, China).

### CUT&RUN analysis

After removing adaptor sequences and low-quality reads by cutadapt (version 4.6), CUT&RUN paired-end reads were aligned to the human genome (GRCh38) using Bowtie 2 (version 2.3.5.1 by Ben Langmead) with a default setting. Samtools (version 1.10) was used to generate bam files and quality controls were done using deepTools. Normalisation factors were calculated using CUT&RUN greenlist following the protocol of (de Mello et al., 2024). We use the deepTools “bamCoverage” with option --scaleFactor to generate scaled bigwig tracks. Tracks were visualised with IGV (version 2.12.3). Mean bigwig files were generated using WigToBigWig (version 2.9). Peak calling was done using SEACR (Meers et al., 2019) with the following parameters: normalization (non), peak calling (stringent), threshold (0.01). Peaks with less than 300 bp of distance were merged using bedtools. Profile plots were generated using deepTools functions computeMatrix and plotProfile. We downloaded from the ENCODE portal (Zhang et al., 2020, https://www.encodeproject.org/) the bigwig files with the following identifiers: ENCFF635GGC (H2AZ), ENCFF446DQZ (H3K27ac), ENCFF232NCT (H3K4me1), ENCFF885ESG (H3K4me3); and the bed files with the following identifiers: ENCFF446DQZ (H3K27ac), ENCFF232NCT (H3K4me1). R (version 4.4.1) with package DiffBind (version 3.14.0) were used to perform differential binding analysis. Peak annotation was performed with CHIPseeker package (version 1.40.0) with a TSS range of a -500, 500 bp window. Over-representation analysis was performed as described above for the RNA-seq.

### Statistical analyses and figures

Histograms and statistical analyses were performed using R. Each replicate corresponds to MuSC derived from a different HC or DM patient. To perform Student t-test or Anova, we verified normal distribution of samples using Shapiro test and variance equality with Fisher test. Wilcoxon or Mann-Whitney u-test was applied in absence of normality for the sample distribution. P-Values are depicted on graphs as follows: *<0.05; **<0.01; ***<0.001; ****<0.0001. Biorender.com was used to generate figures, schemes and models.

### Antibodies

All the primary and secondary antibodies used in this study, together with the species, the references and the dilutions for immunofluorescence and western blotting, are summarized in the Key Resources Supplementary Table 7 in Appendix.

## Data access

The RNA-seq and CUT&RUN datasets have been deposited in the Gene Expression Omnibus (GEO; http://www.ncbi.nlm.nig.gov/geo/) under the accession numbers GSE298208 and GSE298210. The secure token to allow review of the GSE298208 record is: **ulolsguwptqbvmx**. The secure token to allow review of the GSE298210 record is: **kfsxcuumllmfzyd**. ChIP-seq datasets of histone PTM in HC-MuSC were retrieved from ENCODE (https://www.encodeproject.org/).

## Competing interest statement

None declared.

## Funding

INMG-PGNM laboratory is funded by grants from the Centre National de la Recherche Scientifique (CNRS), Institut National de la Santé et de la Recherche Médicale (Inserm), University Claude Bernard Lyon 1, and AFM-téléthon. This work was supported by the French National Agency for Research-ANR through grants ANR-21-CE17-0018 IFN-Epi-IM to P.L., B.C., O.B., AFM-Téléthon (MyoNeurALP alliance to P.L. and B.C.), ANR-10-LABX-61 LabEX DEVweCAN and ANR-20-COV9-0004-01 to P.L., Ligue contre le cancer and UCBL-SENS to A.C.

## Acknowledgement

We thank William Jarassier for guidance on bioinformatic analyses. P.L. team is member of the Groupements de Recherche, ‘’Dynamique des interactions entre chromatines virale et cellulaire’’ (DYNAVIR, GDR2194), and ‘’Architecture et Dynamique du Noyau & des Génomes’’ (ADN&G) funded by the CNRS. P.L. is a CNRS Research Director.

## Author Contributions

Conceptualization: AC, BC, OB, PL, YA; Methodology and investigation: all authors; Analysis and validation: AC, BC, OB, PL, RM, WB, YA; Resources: LG, BC, OB, RM; Writing - Original Draft: AC, BC, PL, WB; Writing - Review & Editing: all authors; Supervision and funding acquisition: OB, YA, AC, BC, PL

---

**Supplementary figure 1.**
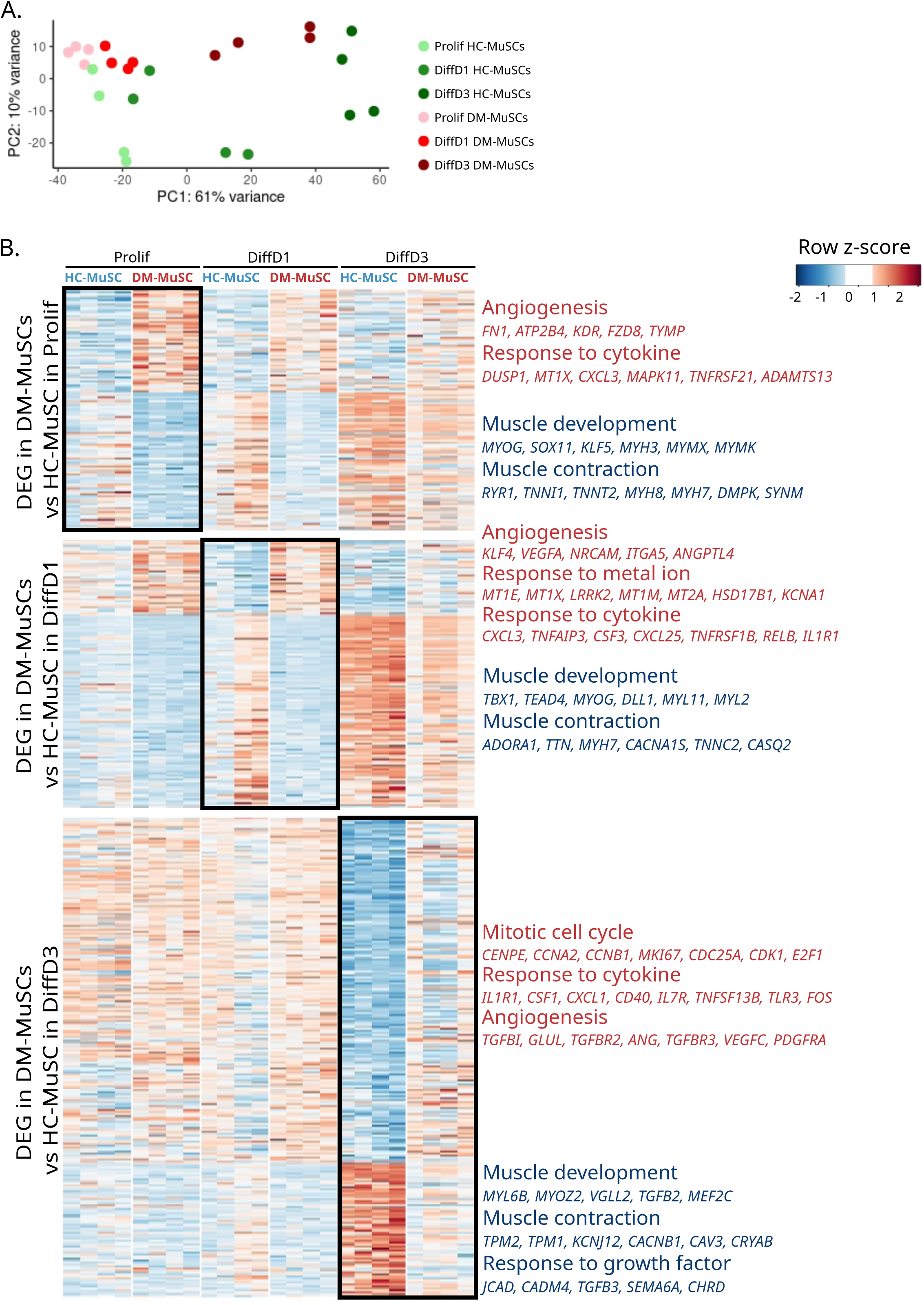
DM-MuSCs have profound transcriptomic defects. (**A**) PCA of gene expression of HC- and DM-MuSCs in Prolif, DiffD1 and DiffD3 conditions. (**B**) Heatmap of the expression of DEGs between DM- and HC-MuSCs in Prolif, DiffD1 and DiffD3 conditions. On the X axis are samples sorted by culture condition and cell type. On the Y axis DEG set associated with genes up-regulated in DM-MuSCs and genes downregulated in HC-MuSCs. Overrepresented GO:Biological Process terms of each DEG set with a selection of representative genes are indicated in red for genes up-regulated in DM-MuSCs and in blue for genes downregulated in HC-MuSCs.

**Supplementary figure 2.**
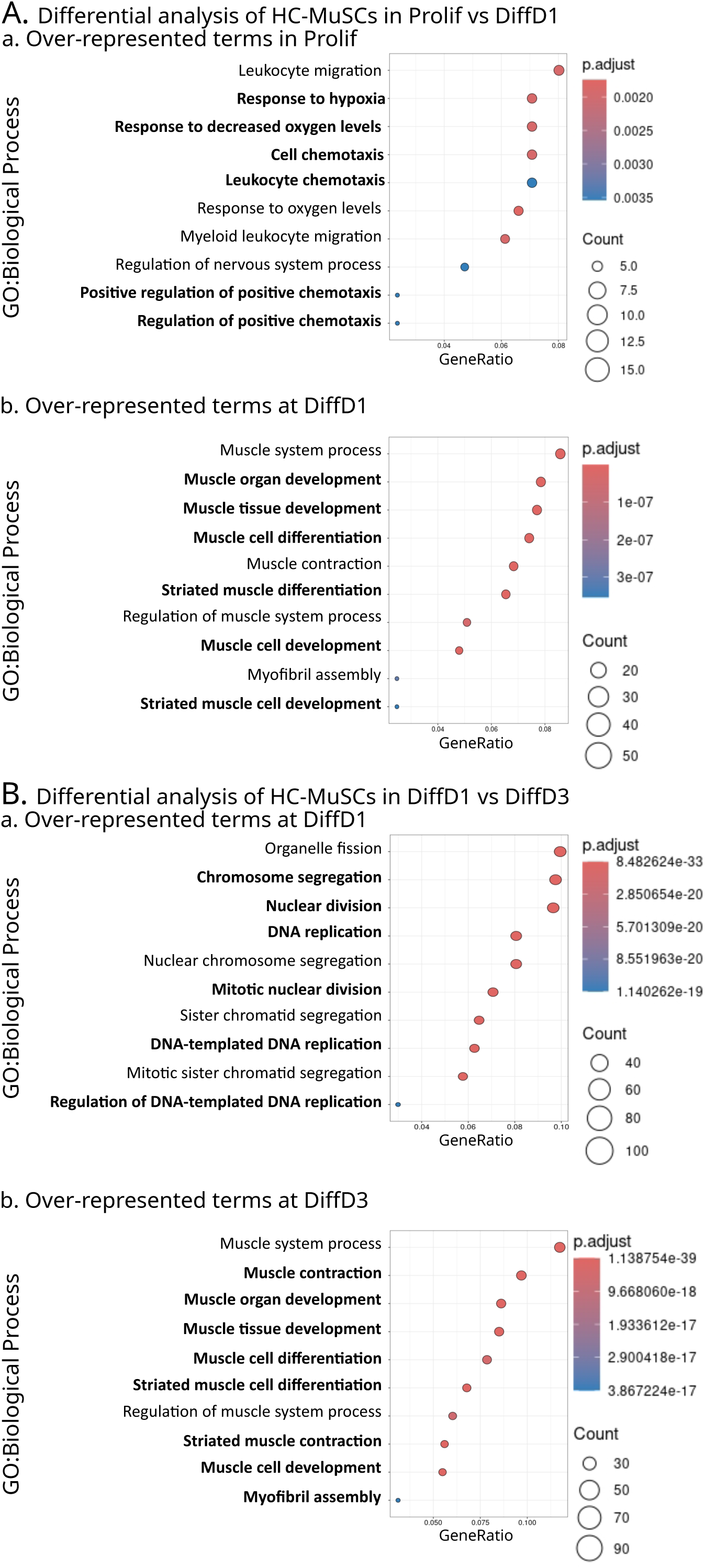
Analysis of differentially expressed genes between HC-MuSCs in proliferation and differentiation. (**A**) to (**B**) Gene ontology biological process (GO:Biological process) over-representation analysis of the DEG between HC-MuSCs in Prolif *vs*. DiffD1 and between HC-MuSC at DiffD1 *vs*. DiffD3. (**Aa**) Over-represented terms in Prolif (Prolif *vs*. DiffD1). (**Ab**) Over-represented terms at DiffD1 (Prolif *vs*. DiffD1). (**Ba**) Over-represented terms at DiffD1 (DiffD1 *vs*. DiffD3). (**Bb**) Over-represented terms at DiffD3 (DiffD1 *vs*. DiffD3).

**Supplementary figure 3.**
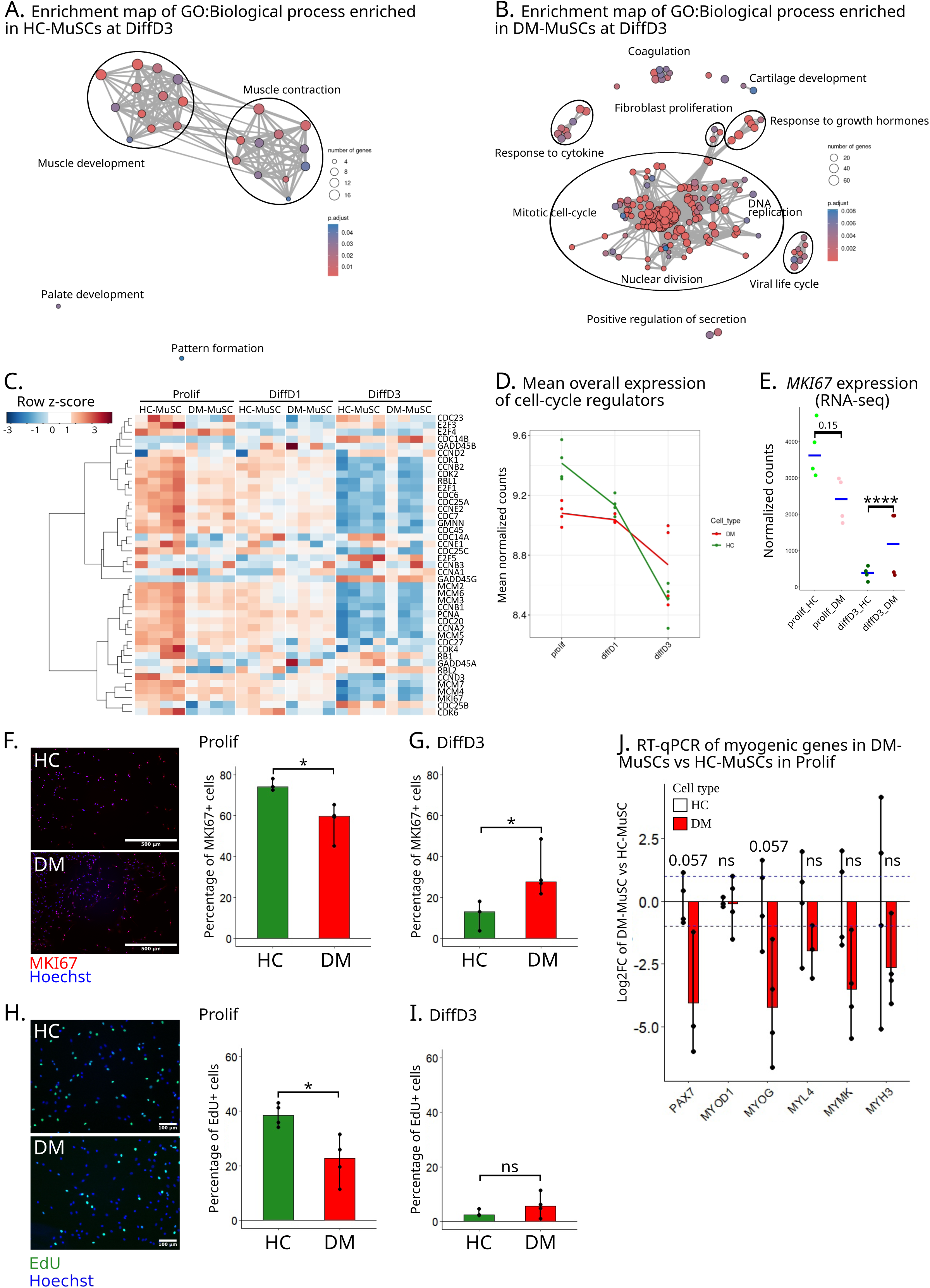
DM-MuSCs show blunted plasticity in response to differentiation stimulus. **(A**)-(**B**) Enrichment map of GO:Biological Process enriched in DEGs between HC- and DM-MuSCs at DiffD3. Cluster of similar terms are encircled and annotated. (**A**) Enrichment map of over-represented GO:Biological Process terms in HC-MuSCs (**A**) and DM-MuSCs (**B**) at DiffD3. () Heatmap of the expression of genes coding for cell-cycle regulators in DM- and HC-MuSCs in Prolif, DiffD1 and DiffD3 conditions. (**D**) Mean expression per condition of all genes coding for cell-cycle regulators between DM- (red) and HC- (green) MuSCs. (**E**) Expression of *MKI67* from the RNA-seq data. Blue crossbars indicate median for each cell type in each condition. P-value significance between HC- and DM-MuSC is indicated on the graph. (**F**)-(**G**) Immunofluorescence for MKI67 (red) in HC- and DM- MuSCs (nuclei blue) and quantification as percentage of positive cells in Prolif (**F**) and Diff3 (**G**) conditions. (**H**) EdU staining (green) of HC- and DM-MuSCs (nuclei blue) and quantification as percentage of positive cells in Prolif (**H**) and Diff3 (**I**) conditions. P-value significance between HC- and DM-MuSC is indicated on the graph. (**J**) Expression of myogenic regulatory genes in DM- *vs*. HC-MuSCs from four different patients each, in the Prolif condition, normalized to their mean expression in HC-MuSCs (Log2FC). P-value significance between HC- and DM-MuSC is indicated on the graph.

**Supplementary figure 4.**
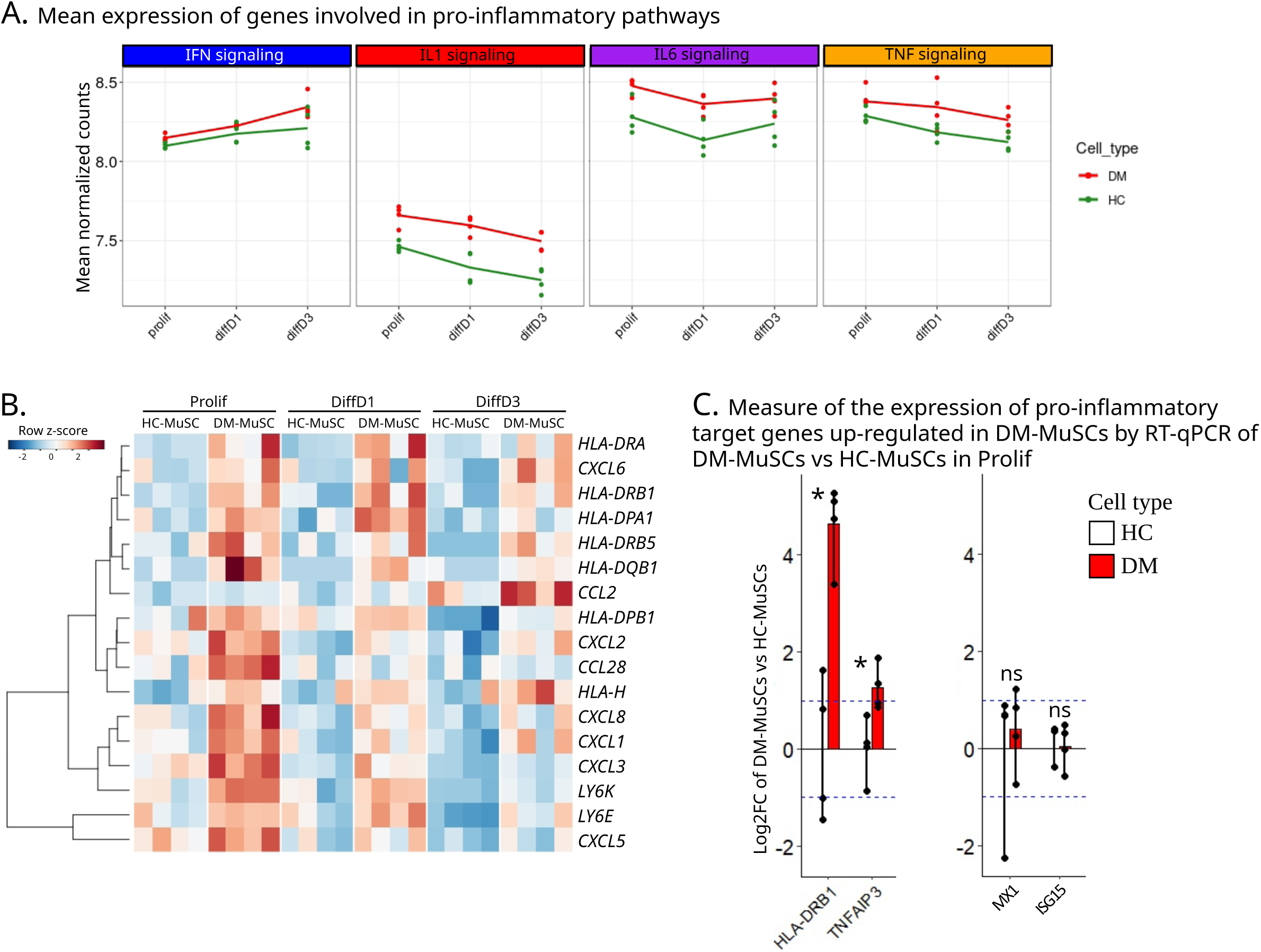
Pro-inflammatory pathways activation in DM-MuSCs. (**A**) Mean expression by cell type and by condition of all the genes involved in type I, II and III IFN signaling (blue panel), IL1 signaling (red panel), IL6 signaling (purple panel) and TNF-⍺ family signaling (orange panel) between DM- (red) and HC-MuSCs (green) in Prolif, DiffD1 and DiffD3 conditions. (**B**) Heatmap of the expression of genes coding for MHC, chemokines and T-cell receptor regulators in DM- and HC-MuSCs in Prolif, DiffD1 and DiffD3 conditions. () RT-qPCR of the log2FC expression of pro-inflammatory pathways target genes in Prolif in DM- MuSCs and HC-MuSCs, from four different patients each, normalized to their mean expression in HC-MuSCs. P-value significance between HC- and DM-MuSC is indicated on the graph.

**Supplementary figure 5.**
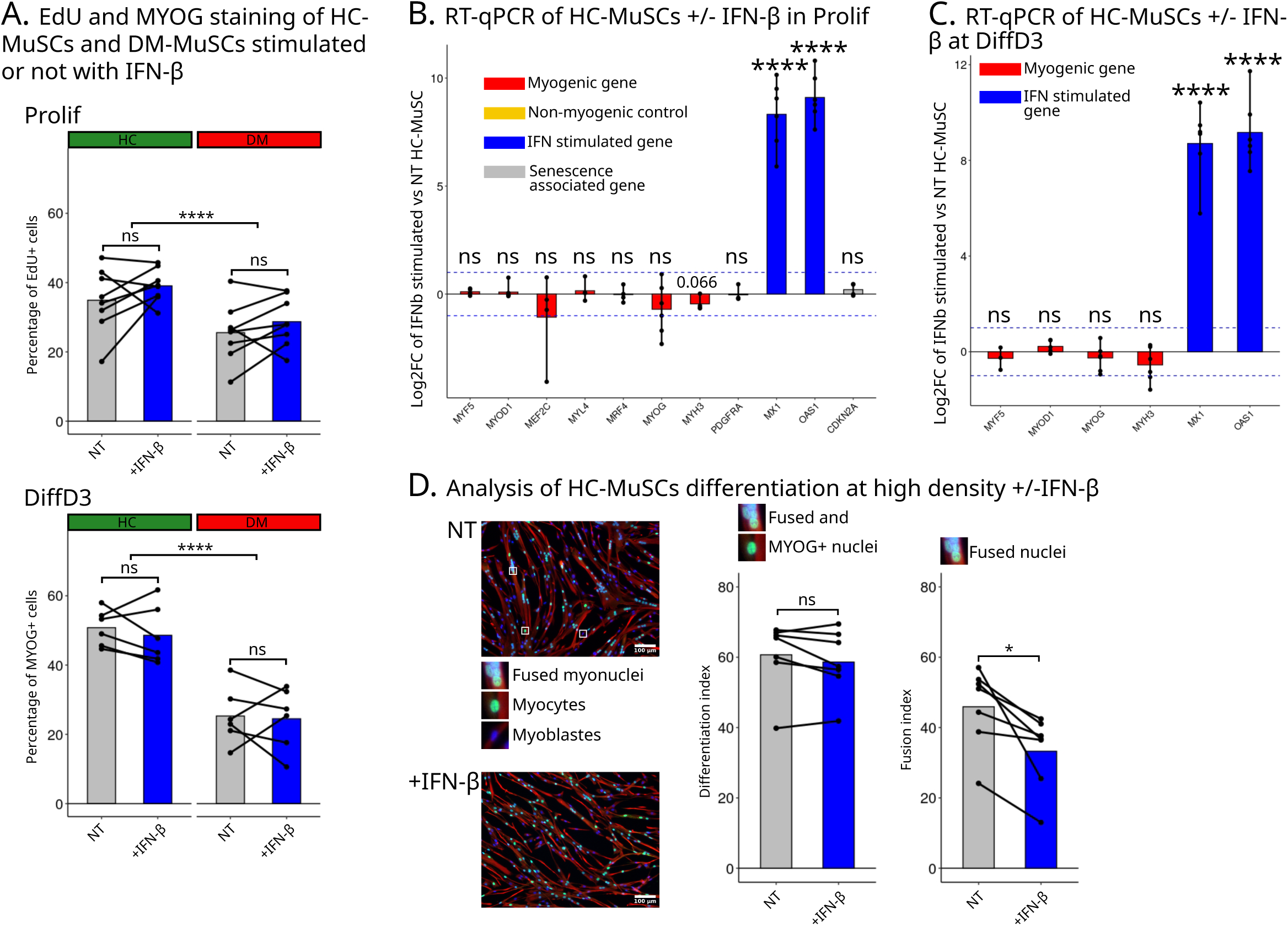
Activation of IFN signaling does not explain the defects observed in DM-MuSCs. (**A**) Quantification of the mean percentage of EdU+ nuclei in HC- and DM-MuSCs from four different patients each, the experience was repeated on the same cells twice at different passages, in Prolif with or without addition of IFN-β (top panel). Quantification of the mean percentage of MYOG+ nuclei in HC- and DM-MuSC from three different patients each, the experience was repeated on the same cells twice at different passages, at DiffD3 with or without addition of IFN-β (bottom panel). (**B**) Expression of myogenic genes, non-myogenic control gene PDGFRA, IFN-stimulated genes and senescence-associated gene CDKN2A in Prolif in IFN-β treated HC-MuSCs from six different patients normalized to their mean expression in untreated HC-MuSCs (RT-qPCR). () Expression of myogenic genes and IFN-stimulated genes at DiffD3 in IFN-β treated HC-MuSCs from seven different patients normalized to their mean expression in untreated HC-MuSCs (RT-qPCR). (**D**) HC-MuSCs from seven different patients in differentiation at high density were treated or not with IFN-β and immunofluorescence for MYOG (green) and desmin (red) was performed (nuclei, blue). Representative fused myonuclei, MYOG+ mono-nucleated myocyte and MYOG- myoblast are zoomed. On the middle panel, quantification of the mean percentage of fused myonuclei and MYOG+ mono-nucleated myocytes between untreated and treated HC- MuSCs. On the right panel, quantification of the mean percentage of fused myonuclei between untreated and treated HC-MuSCs.

**Supplementary figure 6.**
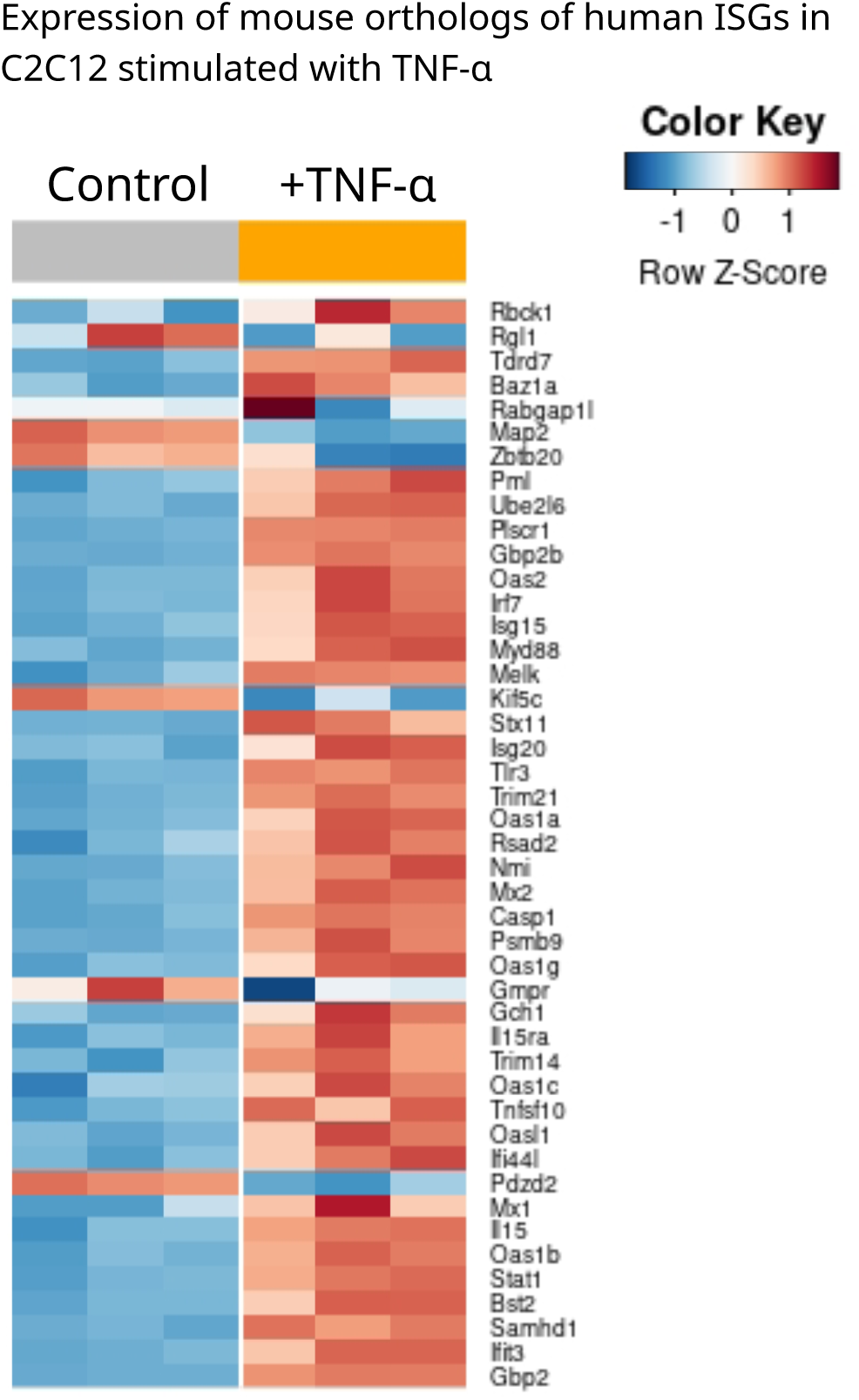
TNF-⍺ stimulation induces the up-regulation of ISG expression. Heatmap of the expression of 45 orthologs of ISGs (McFarlane et al., 2019) in mouse C2C12 cells treated or not with TNF-α in proliferation condition. Data re-analyzed from (O’Brien et al., 2020).

**Supplementary figure 7.**
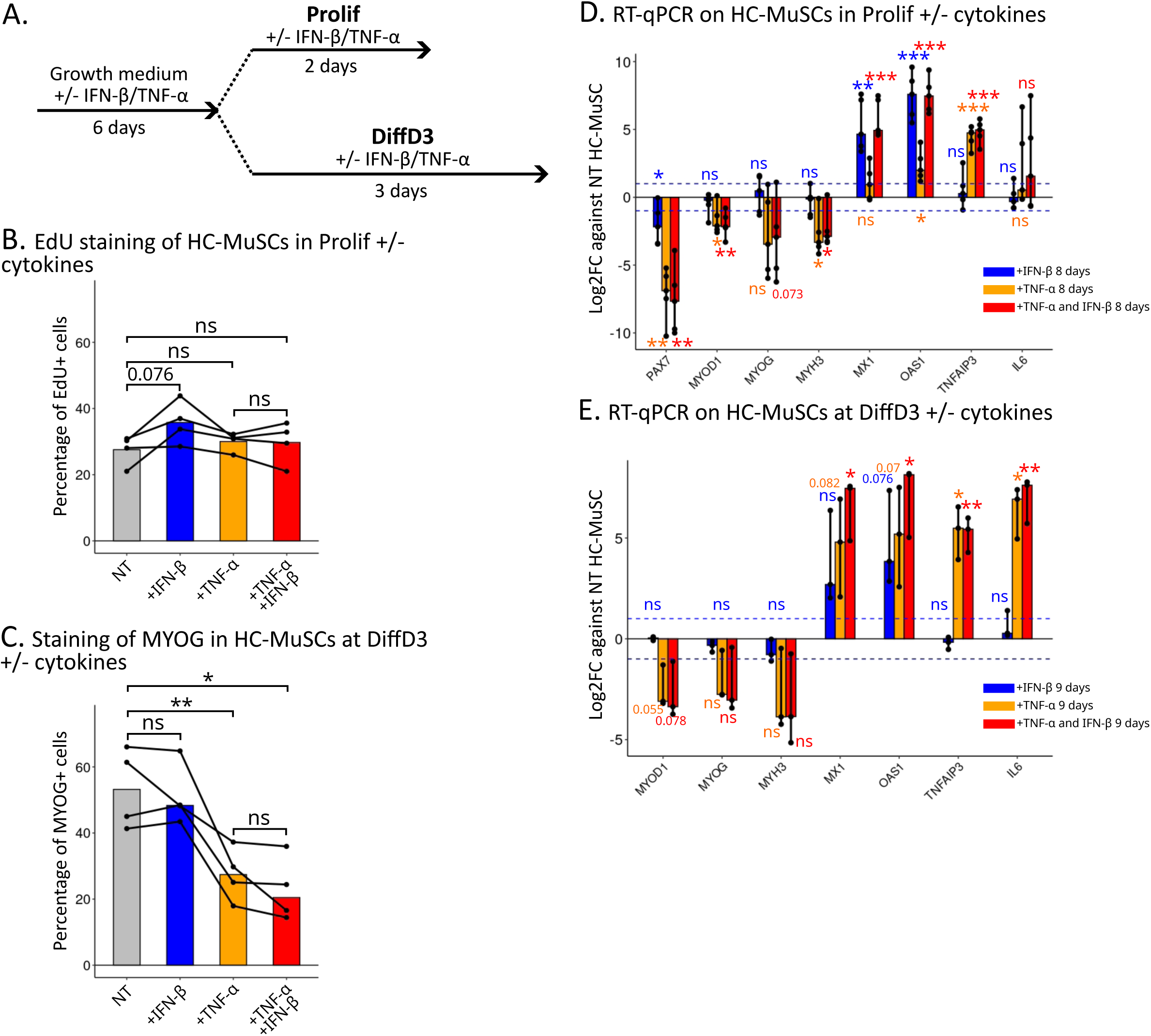
Mid-term cytokine stimulation of HC-MuSCs does not phenocopy patient DM-derived MuSCs. (**A**) Schematic of the mid-term cytokine treatment of HC-MuSCs. (**B**) Quantification of the mean percentage of EdU+ nuclei of HC-MuSCs from three different patients cultured for 8 days with or without IFN-β and/or TNF-α in the Prolif condition. () Quantification of the mean percentage of MYOG+ nuclei of HC-MuSCs from four different patients cultured for 9 days with or without IFN-β and/or TNF-α in DiffD3 condition. (**D**) Expression of myogenic genes, IFN- and TNF-⍺ stimulated genes of HC-MuSCs from five different patients cultured for 8 days with or without FN-β and/or TNF-α in the Prolif condition (RT-qPCR). (**E**) Expression of myogenic genes, IFN- and TNF-⍺ stimulated genes of HC- MuSCs from three different patients cultured for 9 days with or without IFN-β and/or TNF-α in the DiffD3 condition (RT-qPCR).

**Supplementary figure 8.**
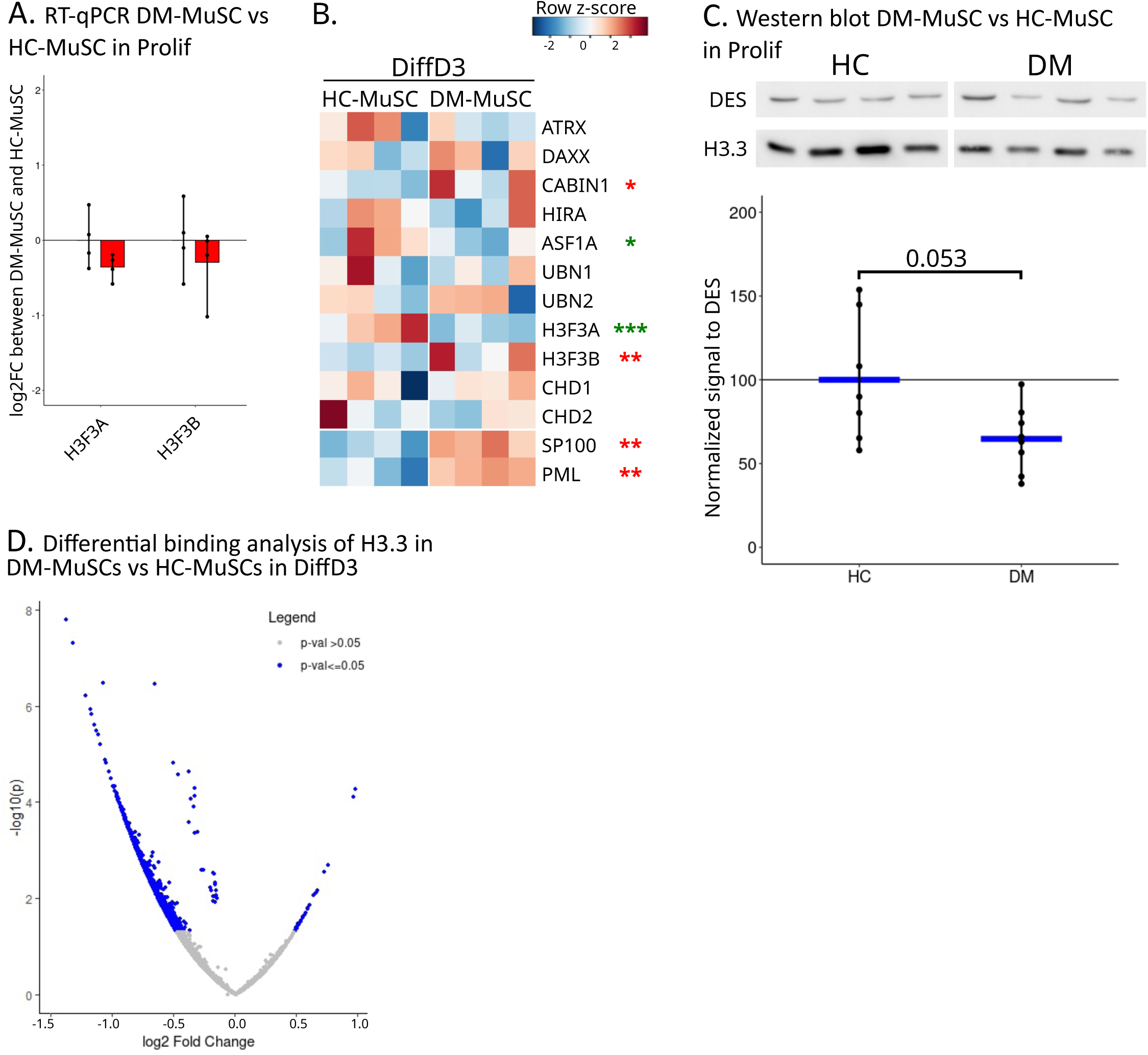
DM-MuSC have a deficit of H3.3. (**A**) Expression of *H3F3A* and *H3F3B* in DM-MuSCs and HC-MuSCs from four different patients each normalized to their mean expression in HC-MuSCs in the Prolif condition (RT-qPCR). (**B**) Heatmap of the expression of H3.3 coding genes, H3.3 chaperones, PML and SP100 in HC- and DM-MuSCs in the DiffD3 condition. Genes down or up-regulated between HC- and DM-MuSCs are labelled with red and green asterix, respectively (the number of asterisks refers to the P value). () Western blot for H3.3 and desmin (DES) in HC- and DM-MuSCs from four different patients each, the experience was repeated on the same cells twice at different passages, with the quantification of the signal of H3.3 normalized to DES for each sample. Blue crossbars indicate mean signal for each cell type. (**D**) Volcano plot of the differential binding analysis of H3.3 CUT&RUN between DM- and HC-MuSCs in the DiffD3 condition. Peaks with a significant differential binding of H3.3 appear in blue.

**Supplementary figure 9.**
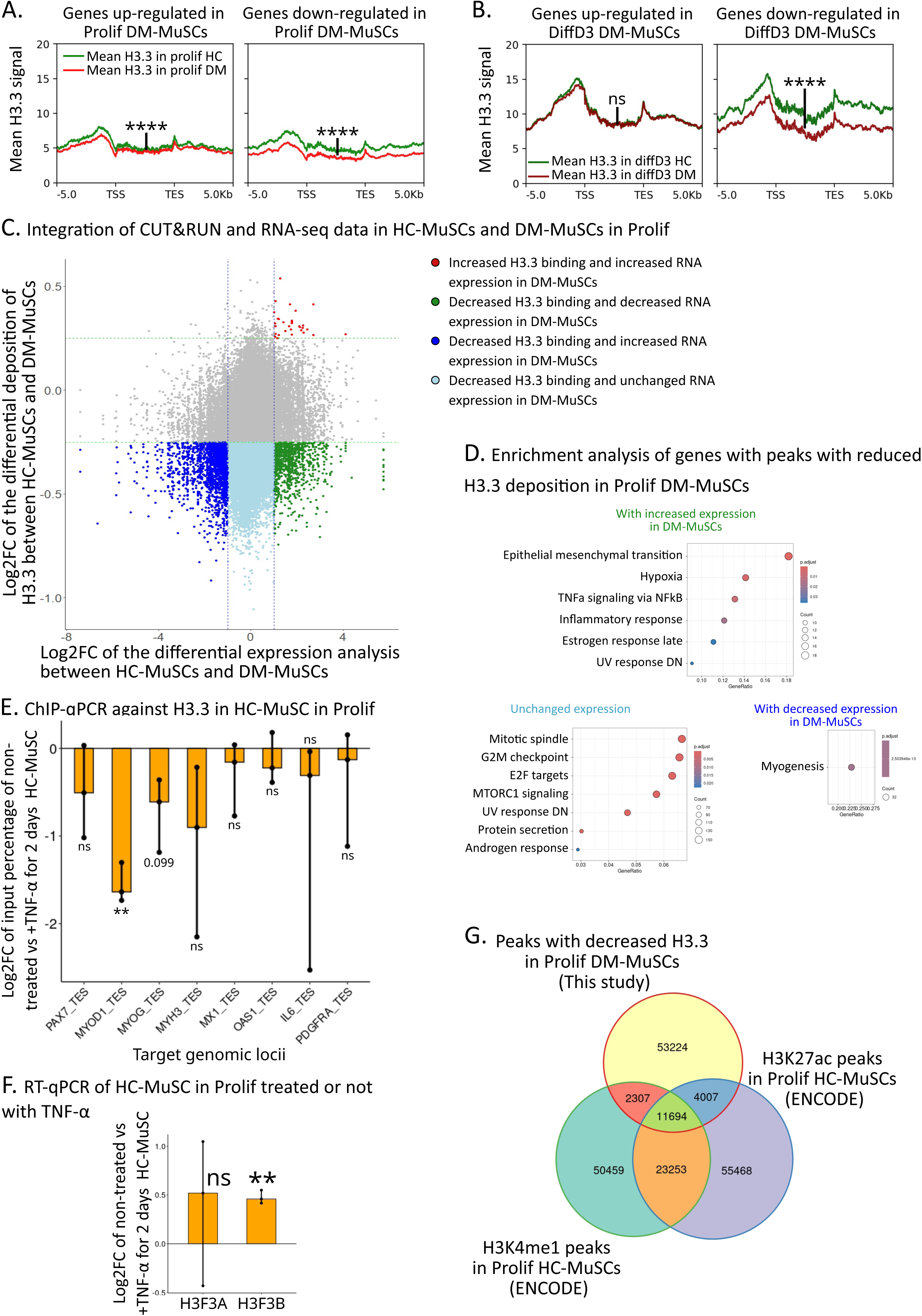
Inflammation driven loss of H3.3 is concentrated on MRFs and enhancers. (**A**) Intensity plot on DEG up-regulated (left panel) or down-regulated (right panel) in DM-MuSCs in Prolif of the mean H3.3 signal in proliferating DM-MuSCs (in red) and HC-MuSCs (in green). (**B**) Intensity plot on DEG up-regulated (left panel) or down-regulated (right panel) in DM-MuSCs at DiffD3 of the mean H3.3 signal in DM-MuSCs (in dark red) and HC-MuSCs (in dark green) at DiffD3. For each panel in (**A**) and (**B**), differential mean H3.3 signal within the -5/+5kb of the TSS and TES of the selected genes was evaluated between HC- and DM-MuSCs, with each gene of the selected panel counted as a replicate. P-value significance between HC- and DM-MuSC is indicated on the graph. (**C**) Integration of the differential binding of H3.3 and differential expression analyses between DM- and HC-MuSCs in the Prolif condition. In x-axis log2FC of the differential expression of genes. In y-axis log2FC of the differential binding of H3.3 in peaks located in genes. (**D**) MSigDB Hallmarks over-representation analysis of genes bound with H3.3 and showing a decreased H3.3 deposition in DM-MuSCs. Left, middle and right panel show genes that with increased, unchanged or decreased expression in DM-MuSCs as compared with HC-MuSCs in the Prolif condition. (**E**) ChIP-qPCR for H3.3 of HC-MuSCs from three different patients treated with or without TNF-α in the Prolif condition. Results in log2FC of the input percentage of H3.3 in the transcription end sites (TES) in TNF-α stimulated HC-MuSCs normalized to untreated HC-MuSCs. (**F**) Expression of *H3F3A* and *H3F3B* of HC-MuSCs from three different patients cultured with or without TNF-α in the Prolif condition (RT-qPCR). (**G**) Overlap of peaks with decreased H3.3 signal in DM-MuSCs, H3K4me1 peaks in HC-MuSCs (dataset from ENCODE) and H3K27ac peaks in HC-MuSCs (datasets from ENCODE, Zhang et al., 2020, https://www.encodeproject.org/).

## Notes

### Competing Interest Statement

The authors have declared no competing interest.

